# A non-canonical feedforward pathway for computing odor identity

**DOI:** 10.1101/2020.09.28.317248

**Authors:** Honggoo Chae, Arkarup Banerjee, Dinu F. Albeanu

**Affiliations:** Cold Spring Harbor Laboratory, Cold Spring Harbor, NY; Cold Spring Harbor Laboratory School for Biological Sciences, Cold Spring Harbor, NY; New York University Medical Center, New York, NY

**Keywords:** mitral and tufted cells, piriform cortex, anterior olfactory nucleus, cortical feedback, concentration invariant odor identity decoding, two photon calcium imaging, PCA, dPCA, linear and non-linear decoders

## Abstract

Elucidating neural circuits that enable robust odor identification remains a fundamental challenge in olfaction. The current leading model states that odor identity is computed within the piriform cortex (PC), drawing from mitral cell input from the olfactory bulb. Here we find that tufted cells, the other principal cell-type of the bulb, which strongly innervate the anterior olfactory nucleus (AON) instead, substantially outperform mitral cells in decoding both odor identity and intensity, acting in a largely feedforward fashion. Cortical feedback from PC specifically restructures mitral cell responses, while feedback from AON preferentially controls the gain of tufted cell odor representations, matching biases in feedforward connectivity. Leveraging cell-type specific analyses, we identify a non-canonical feedforward pathway for odor recognition and discrimination mediated by the tufted cells, and propose that bulb target areas, other than the piriform cortex, such as AON and olfactory striatum, are well-positioned to compute odor identity.

## Introduction

No two stimuli are ever the same. How can we then recognize a face or identify the smell of our favorite bread? The brain readily recognizes different objects in the environment, under widely varying conditions. For example, humans categorize one friend’s face as distinct from another, generalizing across differences in viewing angles, brightness or orientation^1^. The past decade has seen considerable progress in our understanding of the neural mechanisms underlying visual object recognition^1–3^. In parallel, and often synergistically, artificial deep neural networks employing both feedforward and recurrent architectures have been shown to recapitulate core features of sensory transformations along the ventral visual stream^4,5^. Despite recent advances^6–10^, how analogous computations on odors are supported by specific olfactory neural circuits remains an open question.

Two key features of odorants are their identity and intensity. Coffee smells distinct from cheese across most concentrations (proportional to the perceived intensity)^11,12^. Across olfactory behaviors, the brain extracts and acts upon complementary pieces of sensory information. For example, identifying a particular variety of *brie* at a party at varying distance from source requires categorization of the odorants present in the sensory scene independent of their concentration. However, navigating towards the odor source also necessitates estimating the relative concentration of the stimulus.

Odorants are sensed in the olfactory epithelium by olfactory sensory neurons (OSNs) whose axons project to the surface of the olfactory bulb (OB), forming glomeruli sorted by odorant receptor type^13,14^. Glomerular OSN responses are normalized and de-correlated by local inhibitory circuits within the OB^14–18^, and further relayed to higher brain regions in mammals by two distinct populations of output neurons, mitral cells (MC) and, the much less studied tufted (TC) cells, which differ in their size, location, intrinsic excitability, local wiring and activity^19–30^. Tufted and mitral cells project to approximately a dozen brain regions, of which three-layered paleocortical structures such as anterior olfactory nucleus (AON, also known as anterior olfactory cortex) and piriform cortex (PC) are the major targets^31^. Tufted cell projections are biased towards the AON and the olfactory tubercle (OT, olfactory striatum)^20,32,33^. In contrast, while mitral cells project widely and strongly innervate the PC, they send relatively little input to AON^34,35^. In turn, AON and the anterior piriform cortex (APC), as well as most other OB target areas, send numerous feedback axons which primarily target inhibitory bulbar interneurons in different layers^24,36–38^. Cortical-bulbar feedback has been proposed to enable separation of odor representations, sparsening mitral cell responses by targeting specific sets of granule cells which, in turn, inhibit the mitral cells^24,39,40^. However, despite overwhelming evidence for massive top-down projections^41–51^, the specificity and logic of interplay between feedforward and feedback signals between early sensory processing areas and the cortex remain poorly understood.

Over the past decades, computational models as well as experimental results^6–8,52–58^ have proposed that intensity invariant odor representations first emerge in PC, drawing specifically from mitral cell input^6–9,52,54^. In contrast, the AON is thought to estimate the location of odor sources by computing relative stimulus concentration^59,60^ and to process social cues^61,62^. Surprisingly, the role of the tufted cells, the other OB output cell-type, which project strongly to AON, but largely avoid the PC, in odor discrimination and generalization across concentrations has been overlooked. This was partly due to technical limitations of unambiguously identifying mitral versus tufted cells using extracellular recordings.

In this study, we recorded from distinct mitral and tufted cell populations using multiphoton microscopy in awake mice. We asked three specific questions. Do mitral and tufted cells differ in their ability to convey odor identity information to their preferred cortical targets (PC versus AON)? Is cortical feedback from PC and AON specific in controlling the activity of their dominant inputs (mitral versus tufted cells)? What is the relative contribution of feedforward versus feedback input in computing odor identity and intensity? We find that cortical feedback from PC and AON preferentially regulates mitral versus tufted cells respectively, matching biases in feedforward connectivity, and performes different roles. Surprisingly, tufted cells, acting largely in a feedforward fashion, substantially outperform mitral cells in decoding odor identity and concentration. These results, implicate the non-canonical tufted cell-to-AON/OT pathways in mediating odor recognition and discrimination, and re-define our understanding of how and where odor identity and intensity are computed.

### Decoding odor identity with concentration invariance from the olfactory bulb outputs

We investigated whether odor identity and concentration can be decoded from mitral and tufted cell populations, and the extent to which this depends on cortical feedback. Towards this end, we recorded OB output activity in awake head-fixed mice in response to five odorants whose concentration was varied across three orders of magnitude (see Methods, **Extended Data Fig. 1a-f**). Two-photon imaging of GCaMP3/6f^63^ signals enabled us to distinguish mitral versus tufted cells based on the location of their cell bodies in the OB (**Fig. 1a, Extended Data Figs. 1, 5a,b**, see Methods).

**Figure 1:**
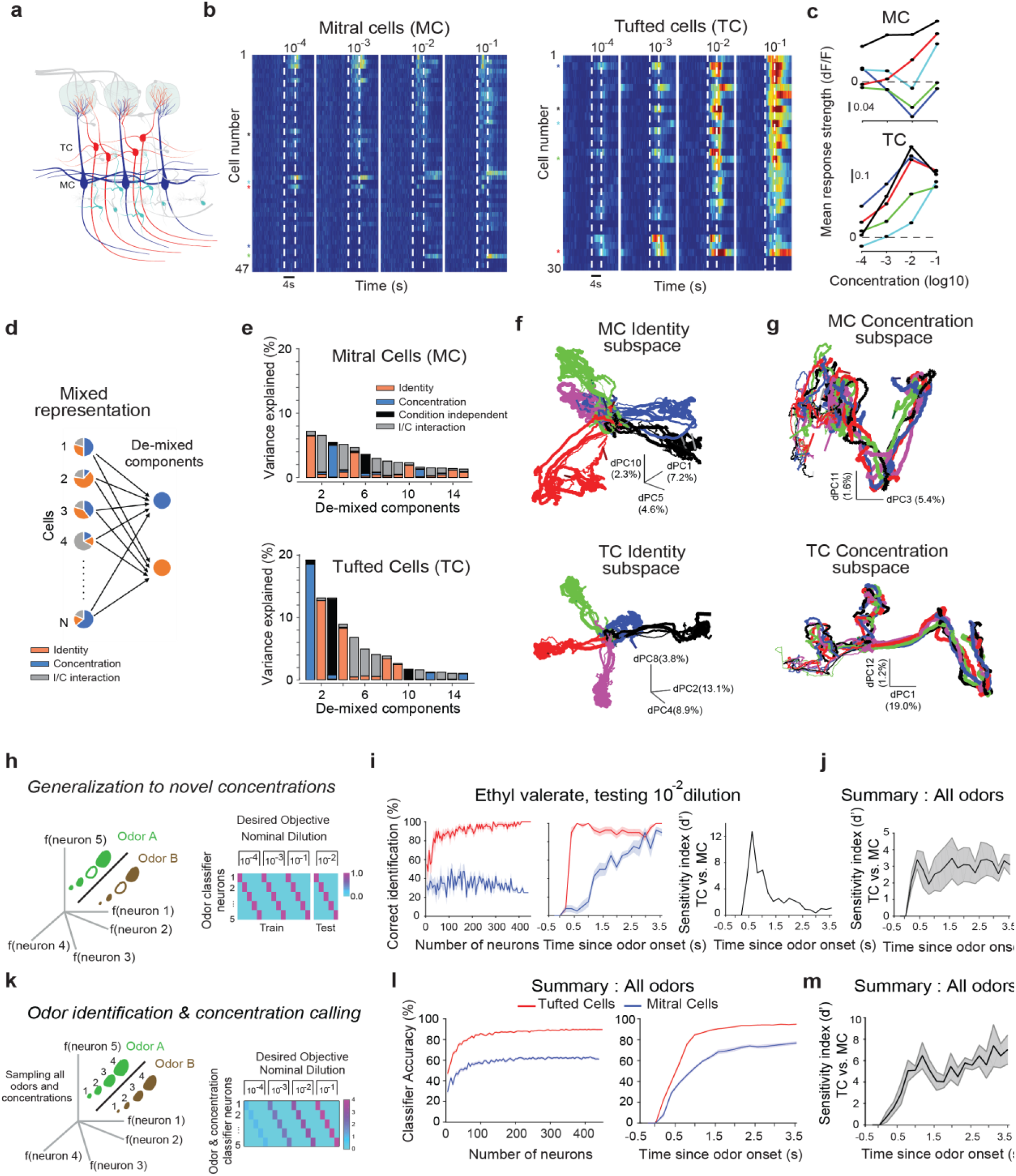
Decoding odor identity with concentration invariance from the olfactory bulb outputs. **a.** Cartoon depicting the two OB output cell-types, mitral cells (MC, blue) and tufted cells (TC, red), located at different depths from pia surface. **b.** (*Left*) Mean peri-stimulus time histogram of simultaneously recorded MC (n = 47, *Left*) and TC (n=30, *Right*) from two example fields of view to increasing concentrations of valeraldehyde. Color indicates normalized change in fluorescence with respect to pre-odor baseline (dF/F0). Dotted lines mark odor presentation (4s). **c.** Mean concentration responses of five example MC (*Top*) and TC (*Bottom*) indicated by colored fiduciary marks in **b. d.** Cartoon showing that mixed representation of odor identity (orange) and concentration (blue) signals in individual neurons can be linearly ‘de-mixed’ as low-dimensional components of the population activity. **e.** For mitral (*Top*) and tufted (*Bottom*) cell ensembles, variance explained by the top 15 principal components identified using demixed-PCA is decomposed into four categories: ‘odor identity’, ‘odor concentration’, ‘interaction between identity and concentration’ and ‘condition independent’ (see Methods); n = 447 MCs and 458 TCs; stimuli: 5 odors, 4 concentrations. **f.** Population trajectories in the neural state space defined by the top three identity principal components for mitral (*Top*) and tufted (*Bottom*) cells. Different colors denote different odorants, while increasing thickness indicates increasing concentration. Total variance explained by the top 3 identity principal components: MC: 14.1%; TC: 25.8%. **g.** Same as **f**, except that neural trajectories are depicted in the sub-space defined by the top two concentration principal components. Total variance explained by the top 2 concentration components: MC: 7%; TC: 20.2%. **h.** *Generalization across concentrations*. (*Left*) The SVM decoder learns to group together any three of four concentrations sampled for a given odorant. Increasing size of odor representations denotes increasing concentration. Cross-validated performance is tested on the ability to classify the fourth concentration previously not used for training (empty circles). (*Right*) Set-up of the decoding strategy where hypothetical classifier neurons (one for each odorant) signal the presence (value =1) of their corresponding odorant for all four sampled concentrations, and its absence (value = 0), for all other odorants in the panel. **i.** (*Left*) Cross-validated classification performance of *generalization across concentrations* for an example odor (*ethyl valerate*) with increasing number of mitral cells (*blue*) and tufted cells (*red*) at a fixed time point (t = 1s). (*Center*) Classification performance for the example odor as a function of time for 200 randomly chosen neurons with bootstrap re-sampling. (*Right*) Tufted and mitral cells performance as quantified by the sensitivity index (d’-d-prime, Methods). **j.** Summary of the difference between MC and TC performance (d-prime) averaged across all odorants in the panel. Chance decoder performance is 0 (Methods). **k.** *Odor identification and concentration calling*. The decoder learns to identify both the odor identity, as well as the relative concentration (on a log scale). Cross-validated performance is evaluated across held-out trials. **l.** (*Left*) Classification performance averaged across all five odorants with increasing number of mitral (*blue*) and tufted (*red*) cells at a fixed time point (t = 1s). (*Right*) Classification performance averaged across all five odorants as a function of time for 200 randomly chosen mitral and tufted cells. **m.** Same as **j,** for *Odor identification and concentration calling*. Shaded areas are SEM unless stated otherwise.

Consistent with previous reports^19,21,64^, we found that mitral and tufted cells had distinct responses to the same stimuli. Notably, mitral cell responses were slow, sparse and phasic, and individual neurons had non-monotonic concentration response curves (**Fig. 1b,c**). In contrast, tufted cells showed fast and sustained responses to odorants, whose magnitude increased monotonically with increasing concentration (**Fig. 1b,c**, **Extended Data Fig. 1b,c**). Additionally, tufted cell responses had significantly lower trial-to-trial variability as compared to mitral cells (**Extended Data Fig. 1f**).

We used de-mixed principal component analysis (dPCA)^65–67^ to investigate whether odor identity and concentration information could be linearly separated from the mitral and tufted cell responses. Success in de-mixing the identity and concentration dimensions implies that a linear combination of neuronal activity can ‘decode’ odor identity with concentration invariance, while, at the same time, a different linear combination of the same neural responses can be used to infer stimulus concentration, irrespective of the odor identity (**Fig. 1d**). Overall, tufted ensembles were better at differentiating odor identity invariant of concentration compared to mitral cells (**Fig. 1e,f**). Tufted cells were also superior to mitral cells in segregating different concentrations of odorants (**Fig. 1e,g**). The degree of de-mixing achieved with this method was substantially larger than by maximizing overall variance using principal component analysis (PCA, **Extended Data Fig. 2a-d**, see Methods).

To quantify directly the ability of mitral and tufted cell ensembles to perform odor identification, we used linear (Logistic regression) and non-linear decoding schemes (Support Vector Machines, SVM, Methods). Specifically, we analyzed the performance of mitral and tufted cells in two decoding schemes inspired from analogous computations necessary for solving olfactory behavioral tasks. First, we probed for generalization to a novel concentration, after odor identity was learned from a test set of concentrations (**Fig. 1h-j**, *generalization across concentrations*).

Second, we aimed to decode both odor identity and concentration, wherein each classifier neuron was tasked with identifying the presence of the corresponding odorant, as well as simultaneously reporting its relative concentration (**Fig. 1k-m**, *odor identification and concentration calling*, Methods).

To account for intrinsic differences in response magnitude across mitral and tufted cells, their activity was z-scored before performing any decoding analyses, and found to span the same range (**Extended Data Fig. 1e**, Methods). By systematically varying the number of cells included in analysis, we trained, evaluated and cross-validated the decoders’ performance at different time points from odor onset (SVM decoder performance at chance is 0, see Methods). Tufted cells substantially outperformed mitral cells in odor generalization across concentrations, both in time, and with respect to the number of neurons required for comparable accuracy (earlier and fewer cells, **Fig. 1i,j**). The differences in tufted versus mitral cells performance were robust and appeared early, within behaviorally relevant timescales (200-500ms) from odor onset as quantified using a sensitivity index (d’, Methods, **Fig. 1i,j, Extended Data Fig. 1a-c**). Importantly, small subsets of randomly selected tufted cells (~10s) were sufficient for successful decoding (**Fig. 1i,l**), highlighting the distributed nature of odorant representations. Tufted cells were also superior to mitral cells when specifically trained to assign odor identity irrespective of concentration (*odor recognition*, cross-validated using held-out trials, **Extended Data Fig. 3a-e**, see Methods). Furthermore, tufted cells were better at simultaneously decoding odor concentration, as well as identity compared to mitral cells (**Fig. 1k-m**).

Thus, tufted cell ensembles carry sufficient information to infer odor identity with concentration invariance, as well as to extract relative odor concentration. Taken together with the known biases in the projection patterns of mitral versus tufted cells^20,32–35^, these results indicate that OB target areas, other than the piriform cortex, such as AON and olfactory tubercle, which receive strong tufted cell input, are well-positioned to compute odor identity.

### Cortical feedback preferentially regulates the activity of its dominant OB input

The responses of mitral and tufted cells are shaped both by feedforward input from OSNs, local interactions via interneurons, as well as top-down feedback from the cortex and other brain regions^23,24,33,36–38^. We investigated the specificity of cortical feedback from the PC and AON onto mitral and tufted cells, as a first step towards evaluating how cortical feedback affects the decoding of odor identity and concentration in the OB outputs. To do so, we reversibly silenced activity in AON or APC using muscimol (a GABA-A receptor agonist, **Extended Data Fig. 4a,b**, Methods^24^), and probed the changes in odor responses of mitral and tufted cells. APC sends feedback projections only ipsilaterally, while AON feedback projections run bilaterally^33^ (**Extended Data Fig. 4c,d**). We therefore monitored the change in mitral and tufted cell responses upon inactivation of the ipsilateral APC, ipsilateral AON, as well as contralateral AON (**Fig. 2a,b, Extended Data Fig. 4a-e**). To account for potential non-specific decay in responses over long imaging sessions (before and after muscimol/saline injection), all changes in mitral and tufted cell activity were normalized to saline regression controls (**Extended Data Fig. 5a-c**, Methods).

**Figure 2.**
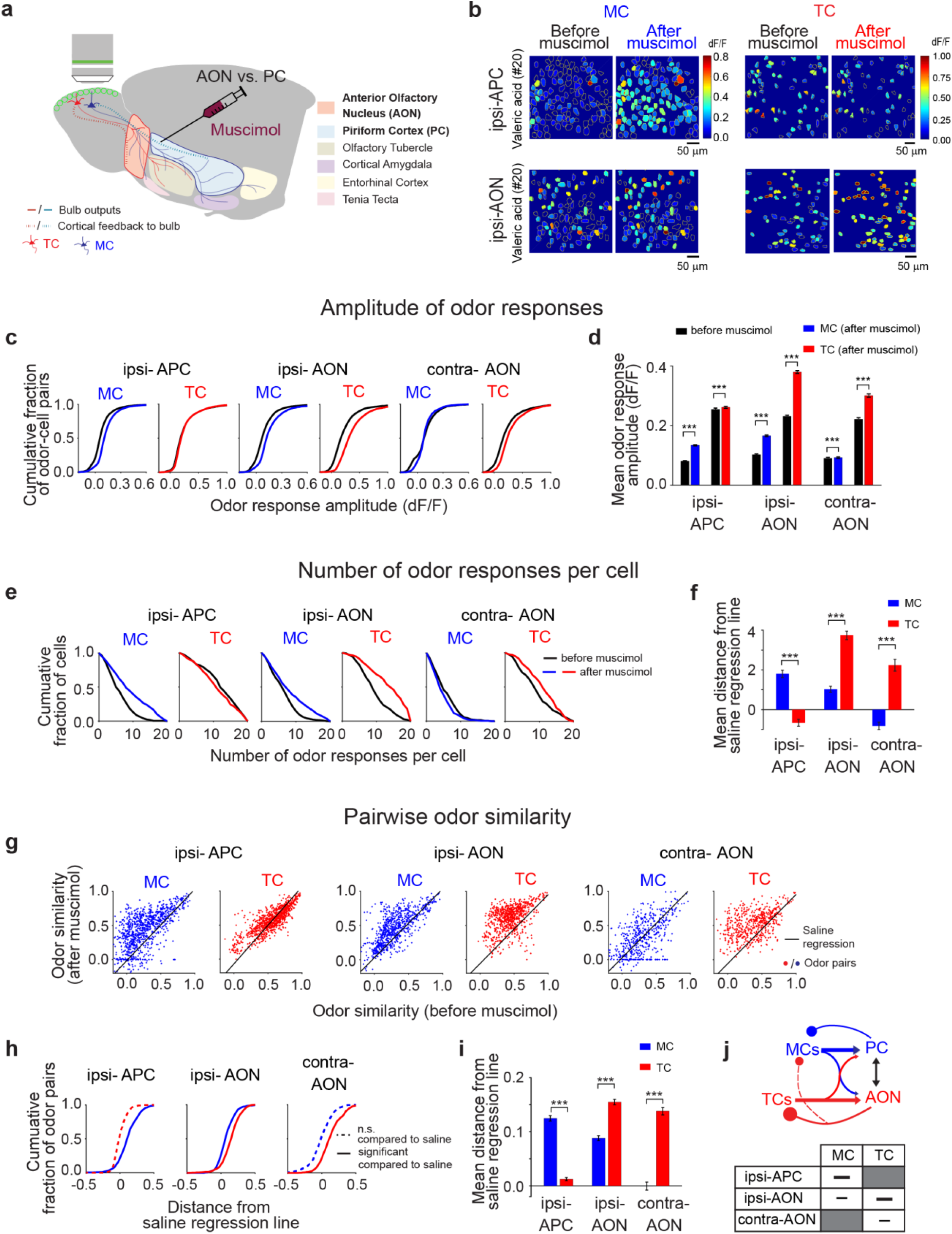
Feedback from the anterior piriform cortex (APC) and anterior olfactory nucleus (AON) to the olfactory bulb differentially regulates the odor responsiveness and pairwise correlations of odor representations in mitral versus tufted cells. **a.** Schematic of experimental procedures: cartoon representation of olfactory bulb (OB) and its major projection target areas including the anterior olfactory nucleus, AON, piriform cortex, PC, olfactory tubercle, cortical amygdala, lateral entorhinal cortex and tenia tecta. Mitral cells (MC, blue) strongly innervate the PC, while tufted cells (TC, red) preferentially project to the anterior olfactory nucleus (AON) and olfactory tubercle. Dotted lines indicate cortical feedback to the bulb from PC and AON. Odor responses of MC and TCs were sampled in awake head-fixed mice via two photon imaging of GCaMP3/6f signals before and after suppression of activity in the anterior piriform cortex (APC) and AON respectively via muscimol injection. **b.** Average odor responses (fluorescence change, dF/F0) of mitral (leftmost two columns) and tufted cells (rightmost two columns) in two example fields of view (220μm and 150μm below the surface) before (*Left*) and after (*Right*) muscimol injection into APC (*Top*, ipsi-APC) and AON (*Bottom*, ipsi-AON). **c.** Cumulative distribution of MC and TC-odor response pairs as function of dF/F0 responses amplitude before (black) and after (blue: MC, red: TC) muscimol injection into ipsi-APC, ipsi-AON and contra-AON. **d.** Summary of mean odor responses amplitude (dF/F0) of MC and TC before (black) and after (blue: MC, red: TC) muscimol injection into ipsi-APC, ipsi-AON and contra-AON. *\*\*\*p<0.001*, One-sided Wilcoxon sign-rank test. **e.** Cumulative distributions of number of odors in the panel that individual mitral and tufted cell responded to before (black) and after (MC: blue, TC: red) muscimol injection into ipsi-APC, ipsi-AON and contra-AON. **f.** Summary of mean distance from saline regression line for changes in the number of odor responses per cell after muscimol (blue: MC, red: TC) into ipsi-APC, ipsi-AON and contra-AON. \*\*\**p<0.001*, Wilcoxon rank-sum test. **g.** Scatter plots of pairwise odor similarity of mitral cells (MC, blue) and tufted cells (TC, red) responses before and after muscimol injection into ipsi-APC, ipsi-AON and contra-AON; each dot represents one pairwise odor-to-odor comparison before and after muscimol injection; combined responses from responsive mitral and tufted cells respectively across all sampled fields of view (ipsi-APC MC: n=950 odor pairs, TC: n=950 odor pairs; ipsi-AON MC: n=760 odor pairs, TC: n=760 odor pairs; contra-AON MC: n=570 odor pairs, TC: n=570 odor pairs; number of odor pair comparisons is summed across the fields of view). **h.** Cumulative plots of distance distributions from saline regression of MC (blue) and TC (red) odor similarity distributions after muscimol injection into ipsi-APC, ipsi-AON and contra-AON. **i.** Summary of mean distance from saline regression line for pairwise odor similarity of MC (blue) and TC (red) representations before and after muscimol injection into ipsi-APC, ipsi-AON and contra-AON. *\*\*\*p<0.001*, Wilcoxon rank-sum test. **j.** Cartoon schematics of functional specificity in two long-range loops that engage mitral cells and the APC, and respectively tufted cells and the AON. Unless stated, hypothesis tests were two-sided.

Consistent with our previous results^24^, suppression of the ipsilateral APC specifically modulated mitral, but not tufted cell responses (**Fig. 2c-i**). Compared to saline injection controls, ipsilateral APC suppression increased mitral cell responsiveness (response amplitude and frequency, **Fig. 2c-f**), as well as pairwise similarity in the mitral cell odor representations (Odor similarity, **Fig. 2g-i**, see Methods). Tufted cell responses were largely unaffected by APC suppression (**Fig. 2c-i**). Conversely, suppression of ipsilateral AON had substantially stronger impact on tufted cells compared to mitral cells. Ipsilateral AON suppression resulted in increased amplitude and number of tufted cell responses and higher pairwise odor similarity (**Fig. 2c-i, Extended Data Fig. 5c-f, 6a,b**). The preferential modulation of tufted cells upon AON suppression was even more apparent upon contralateral AON suppression. Contralateral AON suppression had a negligible effect on mitral cells and specifically boosted individual responsiveness and odor similarity in tufted cells (**Fig. 2c-i, Extended Data Figs. 5c-f, 6a,b**).

Qualitatively, we observed the same results when responses were compared within individual fields of view (**Extended Data Fig. 7a-c**), as well as for matched sample sizes of mitral and tufted cells (**Extended Data Fig. 7d-i**).

Taken together, our results indicate that APC and AON exert preferential suppression and decorrelation of the mitral and tufted cell activity respectively (**Fig. 2j**). This selectivity in feedback regulation of the two OB output channels mirrors the biases in their feedforward connectivity^20,32–35^, thus revealing the existence of two long-range loops which may serve different computations (see Discussion).

### Differential effects of cortical feedback on mitral versus tufted cells

Across sensory modalities, including olfaction, theoretical models suggest that top-down negative feedback precisely balances the strength of feedforward input such as to minimize runaway excitation and/or conveys predictions about upcoming stimuli^39,50,68,71,72^. However, to date, there is little experimental validation of the relationship between feedforward and feedback drives on a cellular level. Our experimental paradigm affords monitoring the same cell-odor pairs with and without cortical feedback. This enabled us to relate the strength and specificity of cortical feedback to the feedforward drive on a cell-by-cell basis (**Fig. 3a-d**).

**Figure 3:**
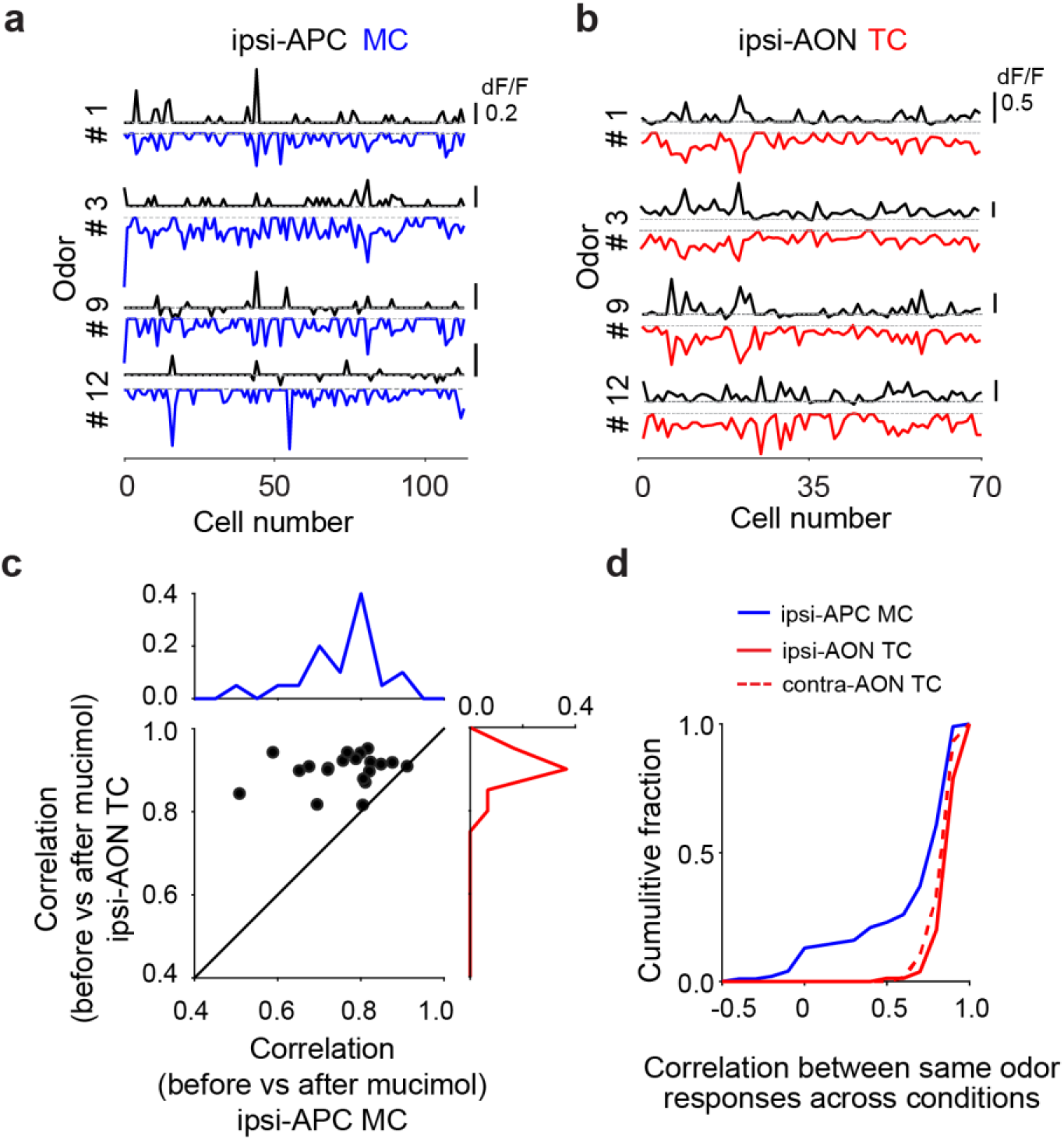
Cortical feedback controls the gain of tufted cell odor representations and restructures mitral cell responses beyond simple scaling. **a.** Mitral cell response spectra from one field of view to example odors before (*upward, black*) and after suppression of APC (*downward, blue*). Non-significant responses were set to zero for visualization purposes (Methods). **b.** Same as **a** for tufted cell response spectra before (*upward, black*) and after suppression of ipsi-AON (*downward, red*). **c.** Scatter plot and histograms for correlation between responses to individual odors before and after ipsi-APC inactivation for mitral ensembles (*blue*) and ipsi-AON inactivation for tufted (*red*) cells. Each data point corresponds to correlation of responses to one odorant before and after muscimol, accrued across FOVs. **d.** Correlation distributions of mitral (blue, MC) and tufted (red, TC) cells responses to individual odors before versus after ipsi-APC (*blue*), ipsi-AON (*red solid line*) or contra-AON (*red dashed line*). In this case, correlation was calculated for each FOV independently.

How does cortical feedback change the ensemble OB output? One possibility is that feedback from APC and AON controls the gain of OB outputs. If this were true, we expect a scaling of odor response amplitudes upon cortical inactivation, while largely preserving the odor tuning of individual neurons. Alternatively, cortical feedback may provide specific information and restructure the population neural activity beyond simple scaling.

APC suppression increased the response probability of mitral cells, rendering them responsive to odors in the panel, which did not evoke a significant response before APC silencing (**Fig. 3a**, *black and blue*). In contrast, odor responses of tufted cells remained self-similar before versus after AON silencing, consistent with a gain control scenario (**Fig. 3b**, *black and red*). To quantify this observation, for each cell type and site of inactivation, we computed the correlation (uncentered correlation coefficient, Methods) between responses to the same odor before and after cortical inactivation. For each odor, tufted cell responses before versus after cortical inactivation (of ipsi- or contra-AON) were significantly more correlated than the mitral cell responses (when suppressing ipsi-APC). This effect was observed both when accumulating responses across fields of view (**Fig. 3c, Extended Data Fig. 8a**), as well as analyzing independently each field-of-view (**Fig. 3d**, Methods). We verified that the effect was specific to the drug condition (**Extended Data Fig. 8b**) and could not be explained by the overall increase in response amplitude after cortical inactivation. For each odor, tufted cell responses remained more self-similar than mitral cells even after after-muscimol mitral and tufted cell responses were scaled down to match the mean of their before-muscimol responses (**Extended Data Fig. 8c-e**, Methods). Similar conclusions were reached when investigating how cortical feedback impacts pairwise odor similarity at the level of mitral vs. tufted cell ensembles (**Fig. 2g-i**) using Receiver Operating Characteristic (ROC) analysis (**Extended Data Fig. 8f-i**).

Taken together, we find that the cortical target of each OB output cell-type predominately controls the activity of its own major input. AON feedback to tufted cells proportionally suppresses the feedforward input drive for each stimulus, thereby regulating the gain of tufted cell odor responses. In contrast, APC feedback restructures mitral cell representations beyond simple scaling^24^.

### Tufted cell ensembles act largely in a feedforward fashion to decode odor identity and concentration

What is the impact of cortical feedback on the ability of OB outputs to decode odor identity and concentration? Since our experiments measure the activity of the same cells before and after cortical suppression, we investigated whether decoding odor identity and/or concentration (**Fig. 1**) requires cortical feedback from APC or AON. For both decoding schemes considered (*generalization across concentrations* and *odor identification and concentration calling*), cortical inactivation reduced the overall decoding performance for both tufted and mitral cells (**Fig.4a-f**). However, the drop in performance was substantially smaller for tufted cells (when inactivating AON) compared with mitral cell based decoders (when inactivating APC), as quantified using both *d’* and a performance difference index (difference between the mean classification performance before versus after cortical inactivation normalized by their sum, see Methods, **Fig. 4c,f**). Similarly, a stronger impact of cortical feedback on mitral compared to tufted cells’ performance was observed when decoders were specifically trained to assign odor identity irrespective of stimulus concentration (*odor recognition*, **Extended Data Fig. 9a-e**). These results indicate that tufted cell ensembles can simultaneously decode odor identity, as well as represent stimulus concentration, using predominantly feedforward processing.

**Figure 4:**
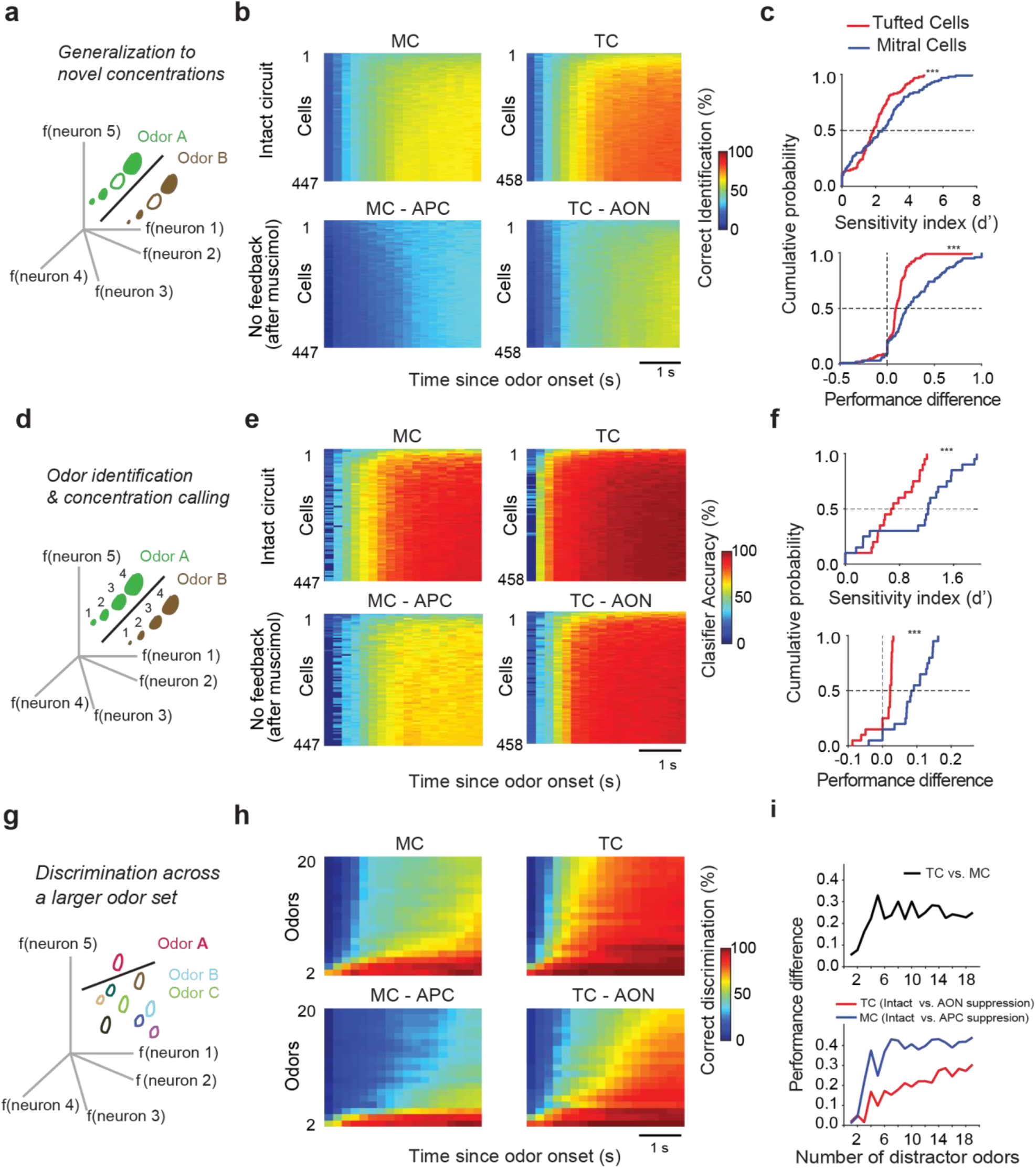
Tufted cell ensembles decode odor identity in largely a feedforward manner and generally outperform mitral cell decoders which depend heavily on cortical feedback. **a**. *Generalization to a novel concentration*. The decoder learns to group together any three of four concentrations sampled for a given odor. Increasing size of odor representation represents increasing concentration. The cross-validated performance is tested on the ability to classify the fourth concentration previously not used for training (empty circles). **b**. 2-D decoder performance map (support vector machine, SVM, non-linear, Methods) as a function of time (abscissa, bin size = 200 ms), while varying the number of neurons included in the analysis (ordinate, bin size = 5 neurons) for mitral cells (*Left*) and tufted cells (*Right*) in the presence (*Top*) and after suppressing feedback (*Bottom*) from the preferred cortical targets (APC for mitral cells and AON for tufted cells). **c.** Summary of the *generalization to novel concentrations* decoding scheme: for both mitral (*blue*) and tufted (*red*) cells, classifier performance difference with and without cortical feedback is quantified using d-prime or a performance difference index (see Methods). In both cases, the performance drop after cortical feedback is significantly higher for mitral than tufted cells. *** indicates p<0.001, paired t-test. **d.** Cartoon schematics for *odor identification and concentration calling* decoding. The decoder learns both odor identity, as well as the relative stimulus concentration (on a log scale). Cross-validated performance is evaluated across held-out trials. Increasing size of odor representation represents increasing concentration. **e.,f.** same as **b** and **c** respectively for *odor identification and concentration calling*. **g.** Cartoon schematics for *discrimination across a larger (20) odor set*. The decoder learns to group increasing number of odors in the panel (1-19) as distinct from the reference odor for a given classification, so as to discriminate any odor in panel from any other. The number of odors included in the analysis is varied systematically from 2 to 20. **h.** 2-D classification performance map (SVM, non-linear) for all four experimental conditions in the *discrimination across a larger (20) odor set* decoding scheme. Abscissa represents the time axis (bin size = 200ms), while the ordinate indicates varying the number of odors included in the analysis using bootstrap re-sampling (ordinate, bin size = 1 odor). **i.** Difference in classifier performance with increasing number of odor distractors. (*Top*) tufted vs. mitral cells (*black line*); (*Bottom*) Intact circuit vs. preferred cortical target feedback suppression for mitral (*blue*) cells and tufted (*red*) cells separately. Chance level performance of the decoder is 0 (Methods).

Animals often sample and discriminate between several odor sources in the same sensory scene. To investigate the performance of mitral and tufted cell ensembles under more naturalistic conditions, we trained classifiers to perform binary odor discriminations of a target odorant from an increasing number (1 to 19) of non-target odorants (see Methods, **Fig. 4g-i).** For discrimination involving only two odorants (one target versus one non-target odor), the ensemble tufted cell decoding accuracy reached 95% within 300-400ms (**Fig. 4h**), matching previous reports of behavioral performance^73^. Increasing the number of non-target odorants led to a gradual drop in tufted cell classification accuracy and increased latency to reach the same performance criterion (**Fig. 4h**). In comparison, mitral cell decoders were slower and fared poorer in discriminating the target odor from non-targets, across the range of discrimination difficulty tested (**Fig. 4h).** The difference in classification performance between mitral and tufted cells increased with progressively difficult binary discriminations (**Fig. 4i**). Moreover, over this range, the relative impact of cortical feedback was substantially higher for mitral cells as compared to tufted cells (**Fig. 4h,i**, *bottom*, **Extended Data Fig. 10**).

Note that the superior decoding performance of tufted cells cannot be attributed simply to their relative higher response amplitude or their lower trial-to-trial variability compared to mitral cell ensembles (**Extended Data Fig. 1b,c**). Cortical suppression increased the response amplitudes of both tufted and mitral cells (**Fig. 2c,d, Extended Data Fig. 5c-e, 7a,d,e, Extended Data Fig. 9f)**. Moreover, the trial-to-trial similarity of mitral responses did not decrease upon APC suppression (**Extended Data Fig. 9i**). However, the decoding accuracy of mitral cell ensembles decreased substantially (**Fig. 4, Extended Data Fig. 10)**. Training classifiers separately on the pre- and post-cortical silencing data sets did not change the advantage of tufted-versus-mitral cell decoders (**Extended Data Fig. 9h**). In addition, the predominantly feedforward performance of tufted cell decoders did not depend upon the specifics of decoders employed, and was consistent across all decoding schemes investigated (**Figs. 1, 4, Extended Data Figs. 4, 9**).

## Discussion

We investigated whether key odorant features such as identity, concentration, as well as concentration invariant identity can be decoded from the two bulb output cell types, and the degree to which the decoding efficiency depends upon cortical feedback from their major targets (PC and AON). Distinct separation of OB outputs into mitral and tufted cells is a feature of land vertebrates, largely absent in fish and amphibians^74^, which appears correlated with the emergence of paleocortex. Over the past decades, a rich body of experimental work and preeminent computational models have proposed that decoding of odor identity (independent or not of intensity) is a central function of the piriform cortex which is strongly innervated by mitral cells^7–9,52^ The recurrent architecture of PC is thought to sculpt the mitral cell input so as to generate concentration-invariant odor identity representations, absent in these OB outputs^6,8^. In agreement with these models, we report that mitral cells are not particularly well-suited for decoding odor identity (**Figs. 1, 4**). However, surprisingly, we find that the other OB output channel, the tufted cells, readily conveys odor identity, as well as odor concentration information in a largely feedforward manner, while feedback from AON controls the gain of their responses to prevent runaway excitation (**Figs. 1, 3, 4**). Importantly, these results were obtained sampling a wider range of concentration and larger odor sets compared to previous studies^7^. Further, our results reflect a conservative estimate of tufted cells’ decoding performance, since in our analysis we considered a 1:1 ratio of tufted-to-mitral cells, while recent anatomical reports estimate this ratio to be as high as to 4:1^75^. As tufted cells innervate mainly brain regions other than PC, such as AON and olfactory striatum, these findings amount to a re-definition of our understanding of how and where odor identity is computed in the mammalian olfactory system.

Thus, decoding of concentration invariant odor identity may not occur solely within the piriform cortex. For example, the feedforward tufted cells to AON pathway is ideally positioned for fast computing of odor identity and concentration during various olfactory behaviors^76^ (**Figs. 1,4, Extended Data Fig. 1c**). These observations are consistent with the proposed role of AON in olfactory search behaviors, as comparator of differences in stimulus intensity across nostrils^59,60^, but also further suggest that AON, and possibly OT, play key roles in representing odor identity^77^. Indeed, comparing with previous behavioral experiments, we note that the decoding performance of tufted cells, rather than mitral cells best matches the behavioral ability of rodents to: 1) extract odor identity in complex odor scenes^73,76,78–80^, 2) identify odors independent of concentration variations^81,82^, and 3) report changes in the concentration of stimuli of interest^11,82,83^. Our findings further raise an intriguing possibility: concentration invariant decoding of anterior piriform cortex responses is inherited from inputs other than mitral cells, either via AON^77^, and/or through direct, sparse tufted cell projections. In summary, while activity from both pathways can be integrated and used for decoding identity, these results indicate that, unexpectedly, the TC-AON pathway plays the major role.

The olfactory circuits involving the OB, AON and APC bear similarity with functional streams in the early visual system. Ganglion cells in the retina project to V1 via the lateral geniculate nucleus in the thalamus, but also send direct input to the superior colliculus^84,85^, which in turn relays to higher cortical visual areas^86^, as well to sub-cortical areas (e.g. peri-aqueductal grey) to support behaviors such as rapid innate visual escape^87^. Similarly, we find that two segregated OB output channels may convey distinct information to the AON and APC. In turn, cortical feedback signals from AON versus APC preferentially target their dominant inputs – the tufted and mitral cells respectively (**Fig. 2**), and appear to perform different computations. A feedforward-feedback loop architecture engaging multiple pathways is powerful, since it enables both implementing different computations on inputs from sensory periphery (i.e. OB, retina, etc.) via parallel streams, as well as cross-talk and comparisons across functional streams. Feedback from AON proportionally suppresses the feedforward input drive for each stimulus, thereby regulating the gain of tufted ensemble activity, without altering the specificity of responses (**Figs. 3, 4**). In contrast, feedback from PC specifically restructures the odor representations of mitral cells (modifies odor tuning), cannot be thoroughly explained by a simple gain control model (**Fig. 3, Extended Data Fig. 8**), and becomes increasingly necessary for hard odor discriminations^88,89^ (**Fig. 4i, Extended Data Fig. 10**). Even in such conditions, tufted cells substantially outperformed mitral cell ensembles in all odor identification schemes analyzed, both before and after cortical activity suppression. Moreover, tufted cells showed lower trial-to-trial response variability, a hallmark of robust stimulus identification, compared to mitral cells (**Extended Data Fig. 1c**), and are more strongly driven by OSN input^30^. Given these findings and, consistent with previous work^52,88–91^, we speculate that the mitral cell-to-PC loop is not primarily involved in the sensory aspects of odor identification, which appear readily supported by the non-canonical TC-to-AON pathway, and rather performs different computations. For example, it may specifically modify odor representations during contextual learning, and/or relay sensory and sensorimotor predictions in complex, fluctuating olfactory scenes. The specificity and functional segregation of cortical feedback action is surprising, given the wide potential for cross-talk either via reciprocal anatomical connections between AON and APC ipsilaterally^92^ or via dedicated tufted and mitral cell specific interneurons in the OB^28^ that may receive convergent inputs from AON and APC. Indeed, our observations that AON feedback, while strongly regulating tufted cell activity also modulates, albeit to a lesser degree, mitral cell responses ipsilaterally, could be due to such cross-interactions.

Our results inform the design of more realistic behavioral tasks^76^ apt to reveal meaningful differences between OB cell-types and the roles of cortical feedback. For example, binary odor discrimination, preferred for its simplicity, may only minimally engage cortical processing, since a handful of tufted or mitral cells with or without cortical feedback seem sufficient to solve it (**Fig. 4**). Future investigation will elucidate the mechanisms by which the TC-to-AON and MC-to-APC loops described here remain specific, the extent of their cross-talk as a function of brain state, and how they support relevant computations for guiding olfactory behaviors.

## Methods

### Surgery

36 adult Tbet-Cre X AI95 or AI38 mice (males and females > 12 weeks old, 25 – 30 g) were administered meloxicam (5mg/kg) and dexamethasone (1mg/kg) 2 hours before surgery. Mice were anesthetized with ketamine/xylazine (initial dose 70/7mg per kg), and supplemented every 45 minutes. Lack of pain reflexes was monitored throughout the procedure. Mice were positioned such that the skull dorsal surface is horizontal, and implanted stereotaxically with cannulae (26 Gauge, Plastics One) bilaterally in the piriform cortex (inserted at 50 degrees from the normal to the brain surface, −4.0 mm (A-P) and 2.4 mm (M-L) from bregma, 7.5 mm deep from surface, corresponding to +1.7 mm A-P, 2.4 mm M-L, 4.0 mm depth from surface at 0 degrees from the normal), and respectively unilaterally in the anterior olfactory nucleus (at 56 degree from the normal, −4.0 mm (A-P) and 1.0 mm (M-L) from bregma, 7.5 mm deep from surface, corresponding to +2.25 mm A-P, 1.0 mm M-L, 4.0 mm depth from surface at 0 degrees from the normal, AON posterior part - AOP). At the same time, a chronic window was implanted above the dorsal aspect of the olfactory bulbs, and a titanium headbar was attached to the skull as previously described^24^ to fixate the animal during the imaging sessions. Meloxicam (5mg/kg) was administered for 5 – 7 days following surgery. Mice were allowed to recover for at least 10 days and further habituated before multiphoton imaging. All animal procedures conformed to NIH guidelines and were approved by the Animal Care and Use Committee of Cold Spring Harbor Laboratory.

### Odor Stimulation

Custom-built odors delivery machines were used to present odors automatically under computer control of solenoid valves^10,24^. Two sets of odors were used: Odor Set A comprising of 5 odors across 4 concentrations, spanning 1:10^4^ to 1:10^1^ nominal oil dilutions, and Odor Set A, comprising of 20 odors, sampled at 1:100 mineral oil dilution, Extended Data Table 1. Odors (1 l/min) were presented in 4s pulses every 1 minute preceded by the acquisition of 10-12s of air baseline and followed by 7-10s of air recovery periods. To minimize odor contamination across trials, during the inter-trial interval, a high flow air stream (>10 l/min) was pushed through teflon coated tubing conduits of the odor delivery machines to an exhaust vent, while the animal’s snout was exposed to fresh air matched at 1 l/min flow rate. Each stimulus was typically repeated 3-5 times before, as well as after-muscimol or saline injections. The concentration of the odors delivered to the mouse for concentration experiments was characterized using a photo-ionization device (PID; Aurora Scientific) and spanned a range between ~0.02% and 10% saturated vapor pressure^16,24^. The same PID was used to determine the time course of the odor waveform and the reliability of odor stimulation. On average, across odors and concentration range sampled, stimuli took 315 +/- 119ms to reach 80% of peak PID value (rising time, **Extended Data Fig. 1a**). This delay in stimulus delivery was accounted for in the decoding analyses (**Figs. 1, 4, Extended Data Figs. 3, 9, 10**) by removing the corresponding frames from the analysis.

### Multiphoton imaging

We used a custom-built multiphoton microscope coupled with Chameleon Ultra II Ti:Sapphire femtosecond pulsed laser (Coherent). The scanning system projected the incident laser beam tuned at 930 nm through a scan lens and tube lens to backfill the aperture of an Olympus 20X, 1.0 NA objective. The shortest possible optical path was used to bring the laser onto a galvanometric mirrors scanning (6215HB, Cambridge Technologies) or resonant scanning head (12 KHz, High Stability 8315K - CRS-12 Set, Cambridge Technologies). Signals were acquired using a GaAsP PMT (H10770PB-40, Hamamatsu), amplified, filtered (DHPCA-100, Femto) and digitized at 200 MHz (NI PXIe-7966R FPGA Module, NI5772 Digitizer Adapter Module). Acquisition and scanning (10 Hz: Odor Set B or 50Hz: Odor Set A) were performed using custom-written software in Labview (National Instruments) including Iris (Keller Lab, FMI). During a typical imaging session, animals were head-fixed under the two photon microscope and habituated to odors and the sound of the scanning galvos (45 min). Laser power was adjusted to minimize bleaching (<40mW). Tufted cells were identified based on the location of their somata in the external plexiform layer (125-175μm from the surface), while mitral cells were identified as a densely packed monolayer of larger somata located 225-275μm deep from bulb surface. Spread across the external plexiform layer (EPL), several subclasses of tufted cells have been described (superficial, middle, internal) at varying depths from bulb surface^75,93^. Here, we probed primarily the activity of middle tufted cells, and further investigation is needed to determine whether different tufted cell subsets relay distinct odor representations.

### Pharmacology

Cannulae were implanted bilaterally for APC and unilaterally for AON suppression experiments (**Extended Data Fig. 4**). Piriform cortex feedback to the OB originates mainly in the anterior apart of the piriform cortex, hence we focused our muscimol suppression experiments on APC^24,37,71,94^. For a given imaging session, muscimol/saline was injected in only one hemisphere. After imaging a given field of view (baseline), muscimol (muscimol hydrobromide, MW=195.01, Sigma) dissolved in cortex buffer was used to suppress neuronal activity in the APC or AON (0.5mg/ml, 1μl injected into APC over 5 min, 0.7 μl injected into AON over 3.5 min). To avoid the spread of muscimol into the olfactory bulb, and accounting for its smaller size, we injected less volume into the AON (0.7 μl) than APC (1 μl). Care was taken to identify the same cell bodies in the field of view before and after the injection of muscimol or saline, waiting 20-30 minutes post-injection before re-starting the imaging session. No apparent changes in animals’ sniffing, whisking or motor behaviors were observed upon muscimol injection. In a previous study^24^, we have quantified the spread of muscimol into APC, calibrating it using comparatively larger volumes of fluorescent muscimol bodipy (Life Science Technologies) to account for the difference in their molecular weights. In this study, we used the same muscimol injection protocol, and identified the injection site (for either muscimol or saline) once the imaging session was completed, by injecting fluorescent muscimol as previously described^24^. Note that any spread of muscimol from AON to APC or vice versa would only result in decreasing the specificity observed in the AON or APC feedback action on the mitral versus tufted cells. Brains were perfused in PFA, and 100-200μm sagittal slices were cut and imaged under an epifluorescence microscope. For control experiments (saline), only cortex buffer was used.

Since AON is a functionally heterogeneous structure comprised of several nuclei^33,95^ which may multiplex information ranging from odor localization and identification to episodic memory^96^ and social cues^61,62^, further studies are necessary to investigate any differences in feedforward-feedback action across different AON nuclei and tufted cell ensembles.

### Tracing of feedback fibers

We checked the distribution of cortical feedback fibers originating in the APC (+1.7 mm A-P, 2.4 mm M-L, 4.0 mm depth from surface at 0 degrees from the vertical), and AON (+2.25 mm A-P, 1.0 mm M-L, 4.0 mm depth from surface at 0 degrees from the vertical, AON posterior part - AOP), post labeling them using targeted viral injections. For visualization purposes, vGlut1-Cre mice were used such as to minimize spurious labeling of migrating (GABAergic) granule cells (and their neuropil) passing in proximity of AON on the way to the bulb. 100nl of AAV2.9-FLOXED-GFP was injected in the AON or APC unilaterally and expression was checked 2 weeks post-infection. 100μm sagittal and coronal bulb slices were obtained after perfusing the brain in PFA. GFP expression was checked under a multiphoton microscope (**Extended Data Fig. 4**).

### Histology

Animals were perfused intra-cardially, the brains preserved in PFA and sliced sagittaly in 100μm thick sections. Slices were mounted on slides using VECTASHIELD Mounting Medium and imaged using an epifluorescence microscope.

### Experimental design

For the concentration-series experiments, 14 mice were employed (APC saline – 4 mice, 4 FOVs; APC muscimol – 3 mice, 4 FOVs; 6 mice for APC saline or muscimol; AON saline – 4 mice, 5 FOVs; AON muscimol – 5 mice, 6 FOVs; 7 mice for AON saline or muscimol). In a subset of the APC suppression experiments, the same animal was imaged more than once. 2 mice (4 FOVs) were injected with saline as well as muscimol in AON, while allowing at least 3 days of intervening recovery time between injections. Each FOV represents a non-overlapping set of mitral or tufted cells. For the larger-odor panel experiments, 22 animals (8 mice for APC; 14 mice for ipsi- and contra-AON experiments) were used. For most experiments, a given brain hemisphere was injected with either saline or muscimol only once. For a subset of experiments (n = 8 mice), both saline and muscimol were injected in the same hemisphere at different times while allowing at least 3 days of intervening recovery time between injections.

## Data analyses

### Pre-processing and detection of significant odor responses

Images were registered laterally (X-Y), and fast Z-movements across individual frames accounted for as previously described^10,24^. Regions of interest (ROIs, mitral and tufted cell somata) were manually selected based on anatomy performed using custom routines in Matlab. To determine the degree of signal contamination by the neighboring neuropil, in a subset of fields of view, for each ROI, a peri-somatic neuropil annulus (10-20μm from the outer edge of the ROI) was generated (**Extended Data Fig. 1d**). Pixels belonging to neighboring non-neuropil ROIs were not included in the annuli. Fluorescence transients were neuropil-corrected as previously described^97,98^ (F_ROI-corrected_ = F_ROI_ - αF⊓_neuropil_). For each ROI, the α parameter was systematically varied between 0 to 1 in 0.25 increments, and the correlation between neuropil-subtracted and raw fluorescence change signals assessed for both z-scored and non-z-scored data. In the z-scored data, neuropil-subtracted signals matched the raw signals with high correlation (slope **~**1) for all α-s considered. In the non-z-scored data, the neuropil-subtracted and raw signals were highly correlated (slope ~1) for α ≤ 0.75 (recent studies used α values ranging from 0.5-0.7^63,97–99^). Thus, neighboring neuropil contamination does not appear to significantly change the odor responses of the sampled mitral and tufted cells. However, given the lack of ground truth for α calling, for the dimensionality reduction and decoding analyses described in **Figs. 1, 4**, and **Extended Data Figs. 2, 3, 9, 10** we used z-scored data without neuropil subtraction.

To determine significance, for each trial and each ROI, we compared the odor evoked normalized fluorescence with values calculated during the air period preceding odor presentations in the session. Responses that exceeded 99.5 percentile of the air period fluorescence distribution, accumulated across all stimuli and repeats for that ROI, were called significantly enhanced as previously described^24^. Responses that were below the 0.5 percentile of the air fluorescence distribution were considered significantly suppressed. An ROI that showed significant responses to an odor in at least two repeats was considered responsive to that odor. Non-significant responses were set to 0 (**Fig. 2e-i, 3a,b, Extended Data Figs. 5f, 6, 7b,c,f-i, 8f-h**). For the analyses performed in **Figs. 1, 3c,d, 4** and **Extended Data Figs. 1, 2, 3, 8a-e, 9, 10**, no thresholding was applied. For **Fig. 2c,d**, and **Extended Data Figs. 5c-e, 7a,d,e**, only response pairs significant in at least one condition (pre- or post-) were used.

#### Odor similarity (odor correlation)

The uncentered odor correlation (odor similarity, *S*^(*A,B*)^) was calculated from the population responses vectors of each pair of odors (A, B) in the panel, in each field of view. Cells responsive to least 2 odors in the panel were included in the similarity analysis.

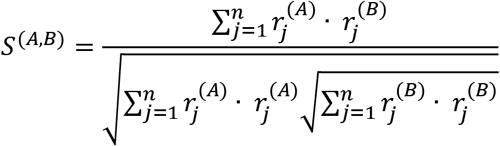

where 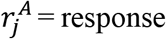 of ROI *r_j_* to odor *A*, 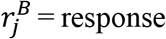 of ROI *r_j_* to odor B, n = number of ROIs.

Same analysis was also performed using Pearson’s correlation, obtaining qualitatively the same results with respect to changes in pairwise odor similarity of mitral versus tufted cell ensembles when suppressing activity in APC versus AON (data not shown).

### Correlation between same odor responses across conditions

The uncentered correlation coefficient was calculated from the population cell response vectors for individual odor across conditions (before- vs. after-muscimol / after-saline injection) accumulated across fields of view (**Fig. 3c, Extended Data Fig. 8a-e**), or for each field of view (**Fig. 3d**). Cells responsive to least two odorants in the panel were included in the correlation analysis. After-injection responses were scaled down (**Extended Data Fig. 8c-e**) such as to match the mean of their before-injection responses. Cells responsive in both conditions, as determined by identifying significant odor responses, were used calculate the scaling down factor. Cell responses to each odor were scaled down independently. Non-significant responses after-injection were left unchanged.

### Dimensionality reduction (PCA and dPCA)

Extracted ROI time courses were assembled in a data cube (N by S by T) of trial averaged dF/F0 responses, where N stands for the total number of neurons included, S is the total number of stimuli and T is the total number of time-bins. To reduce the dimensions of the neuronal population, this data cube was re-shaped into a data matrix (N by ST) and normalized (z-scored) such that each stimulus as a function of time represents a point in an N dimensional neural state space. Neural responses were z-scored to avoid biasing the results to differences in absolute values of response magnitude between the two neuronal classes analysed (mitral versus tufted cells). To find a set of orthogonal directions that maximizes the variance captured from the data, we performed principal component analysis (PCA) and identified the *eigen* vectors of the associated covariance matrix. PCA was performed using built-in ‘princomp’ function in MATLAB. Data projected onto the first three principal components (PCs) is plotted in **Extended Data Fig. 2.** The variance explained by each PC is given by the ratio of its *eigen* value to the sum of all the *eigen* values.

Demixed PCA, a linear dimensionality reduction technique developed by the Machen group^67^ (https://github.com/machenslab/dPCA/tree/master/matlab) was adopted here to decompose the population neural responses into individual components along different features of the odor stimuli. Individual OB output neuron responses multiplex odor identity and concentration representations (**Fig. 1b-d**). Demixed PCA attempts to linearly un-mix these ensemble representations into certain user-defined components, and reveal the dominant neural activity modes (demixed-PCs). In **Fig. 1e-g**, the components were odor identity (I) and odor concentration (C). Data projected onto the first three demixed ‘identity’ PCs or ‘concentration’ PCs are plotted in **Fig. 1e-g.** The same analysis was performed separately for mitral and tufted cells populations. Success in de-mixing odor identity and concentration dimensions implies that a particular linear combination of neurons exists that can ‘decode’ odor identity with concentration invariance, while, at the same time, a different linear combination of the same neural responses can be used to infer the absolute stimulus concentration, irrespective of the odor identity.

### Decoding odor identity (sparse logistic regression and support vector machines)

Odor classification from population neural data was performed using a sparse logistic regression-based decoder with L1 minimization (*lassoglm* function in MATLAB, **Extended Data Fig. 9g**) and a support vector machine *fitcsvm* function in MATLAB, **Fig. 1, 4, Extended Data Fig. 3, 9, 10**) based decoder with either linear or non-linear polynomial kernels. The feature vectors were z-scored mean neuronal responses for each cell in 200ms time-bins. All neurons recorded from multiple field-of-views were pooled, and the number of neurons used in the analyses was varied by sub-sampling (bootstrapping) from this set. The total number of mitral cells (n = 447) and tufted cells (n = 458) were within 2.5% of each other and can be therefore assumed to be approximately 1:1. In all cases, odor classification was analyzed with cross-validation using held-out test dataset. The number of cells considered in the analysis was increased systematically (steps of 5) until all imaged cells well included (**Fig. 1i,l**). For each subset of *k* cells considered, a bootstrap strategy was run 10 times; in each iteration, the decoders were trained and further cross-validated using response from a set of *k* cells was picked randomly with replacement from all cells. Classification performance as a function of time since odor onset (**Fig. 1i,j,l,m**) was evaluated for a fixed number of neurons (n = 200). This procedure was performed 10 times, where 200 neurons were randomly sub-sampled from all neurons recorded. Classification performance as a function of neurons (**Fig 1i,l**) was evaluated at a fixed latency from odor onset (t = 1 s). The SVM was not constrained to pick the best among the alternative possibilities, which would have resulted in chance accuracy of 20% for a five-odor stimulus panel. Instead, failure to accurately classify the corresponding odor identity resulted in a performance accuracy of zero.

The difference between performance distributions across cell-types or pharmacological manipulations, were quantified in two ways - the sensitivity index equivalent to the d-prime (**Figs. 1** and **4**, **Extended Data Fig. 9**) and performance difference index (**Fig. 4, Extended Data Fig. 9**). Sensitivity index (d’), evaluated at each time bin, measured the difference between mean classification performance of the two distributions (m1 and m2) normalized by their standard deviations (σ1 and σ2) as follows:

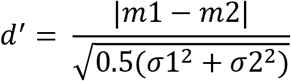

Performance difference index (PDI) was calculated as:

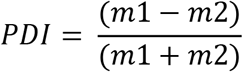

For classification performance comparisons before and after cortical inactivation (**Fig. 4**), decoders were trained on intact circuit neural responses and then tested using neural responses from the *same* neuron after inactivation. In **Extended Data Fig. 9h**, decoders were re-trained after cortical inactivation separately and cross-validated performance was calculated using held-out trial repeats.

Details of the exact type of cross-validation depend on the four different decoding schemes investigated, as described below:

a. *Generalization to novel concentrations*. The decoder learned to group any three of four concentrations sampled for a given odorant together. The cross-validated performance was tested on the ability to classify the fourth concentration previously not used to train. In **Fig. 1i**, training for the lowest two and highest concentrations and testing for the third (second strongest) is shown as example. **Fig. 1l** shows the performance of the decoder in time averaged across all 4 possible combinations of train and test concentrations while varying the number of cells included in the analysis.
b. *Concentration-invariant odor recognition* Neuronal responses to 20 stimuli – 5 odors across 4 different concentrations were used for this scheme and required the classification of all concentrations of each odor as one odor identity. Thus, 100% success would be achieved if each output classifier neuron corresponding to one odorant fired exactly four times, one for each concentration sampled of that odorant, and did not fire in response to any of the other stimuli (**Extended Data Fig. 3**). Cross-validated performance was evaluated by training and testing data-sets, which were taken as different (non-overlapping) repeats of the experimental data (for example, train on 3 repeats and test on the 4th). The plotted decoder performance is averaged across all possible combinations. To test whether the performance of the classifier indeed depended on odor concentrations, for **Extended Data Fig. 3e**, training and testing procedures were identical except concentration labels of each odorant were randomly shuffled for each iteration.
c. *Odor identification & concentration calling* Similar to previous schemes, each odorant has a corresponding classifier neuron. However, each classifier neuron was tasked with identifying the presence of the corresponding odorant (non-zero value), and also simultaneously reporting the relative concentration (on a log scale). The cross-validated performance was evaluated on held-out trials (80% training and 20% testing) same as for *odor recognition*. Classification performance was calculated as a correlation of the decoder output with the objective matrix (**Fig. 1k**).
d. *Discrimination across a larger (20) odor set*. Ensemble mitral and tufted cell responses to 20 diverse odor stimuli were used for this task. For each odor, we considered one output classifier neuron. In the full task, the decoder learns to call the target odor from all other nineteen stimuli. The number of possible odors included in the analysis was varied systematically from 2 to 20 using bootstrap sub-sampling (n=10). Cross-validated performance was evaluated by training and testing data-sets which were taken as different (non-overlapping) repeats of the experimental data. Performance was plotted as average across all possible combinations, while varying the number of odors included (**Fig. 4g-i**). Performance difference index (see above) was calculated as a function of the total number of distractor odorants.

One potential limitation of our study is the slow temporal dynamics of observed responses given the constraints of calcium imaging. We did not investigate fast temporal patterning of spiking in specific sequences with respect to the respiration cycle, which may also contribute to identity decoding^100–103^. However, this is unlikely to affect our main conclusions, since we relied on internal comparisons between the two cell types under identical methodological constraints, and considered the effect of feedback on the same cells before and after pharmacological silencing of the cortex. Further, the decoding differences between mitral and tufted cells reported here represent a lower bound due to the slow calcium dynamics as compared to spiking activity.

#### Quantifying the effect of cortical feedback suppression

Saline regression was performed using all cell-odor pairs (for both response amplitude and odor similarity analyses) before and after saline injection into ipsi-APC or ipsi-AON. Combining all imaged fields of view, a regression line was obtained by minimizing the Euclidian distances from this line to the cell-odor pairs included in the analysis (**Fig. 2f,h,i, Extended Data Figs. 5c-f, 6a,b, 7b,c,f-i,**). A 95% percentile confidence interval with respect to the saline regression line was imposed when calculating the significance of muscimol-induced changes in response amplitude or pairwise similarity. For the response amplitude analysis, only cell-odor pairs showing significant responses in at least in one condition (before or after injection) of saline or muscimol were included.

### Odor similarity-matching number of mitral cells and tufted cells

To match the number of mitral cells and tufted cells used to construct the population responses vectors, we randomly selected 40 cells per field of view, sampling 4 fields of view of mitral cells and 4 fields of view of tufted cells for each iteration of the bootstrap analysis (100 iterations). Saline regression was performed independently for each iteration using the same number of cells and fields of view, and the effects of muscimol injection computed accordingly (**Extended Data Fig. 7**).

### Receiver Operating Characteristic (ROC) analysis

We compared the separation of three distributions (before-, after- and ‘down-scaled’ after-) using Receiver Operating Characteristic (ROC) analysis. For any given threshold value we compared the distribution of odor pairwise similarity values before muscimol injection to that of pairwise similarity values after muscimol injection and respectively of scaled-down after muscimol responses (**Extended Data Fig. 8f,g**). Scaling down was performed to match the average odor response strength of the distribution of pre-muscimol responses. Responses of each odor were scaled down independently. For a given cell-odor pair, only significant responses in either pre or post-muscimol condition were considered. We counted the fraction of odor-odor pairs in the pre-muscimol distribution whose correlation value exceeded threshold (false positives rate) and compared it to the fraction of odor-odor pairs correlation values in the after-muscimol (or scaled-down after muscimol) distribution exceeding threshold (true positives rate). A similar analysis was applied to the saline injection control experiments (**Extended Data Fig. 8h**).

We calculated the percentage of the effect which could be accounted for by gain control as percentage of change in area under the ROC curve (auROC) which can be explained by scaling down after-muscimol odor responses for mitral and tufted cell odor similarity distributions respectively, when suppressing ipsi-APC, ipsi-AON and contra-AON. Since ipsi-APC suppression did not significantly alter TC odor representations, and suppression of contra-AON did not impact MC representations, these table entries were left blank (**Extended Data Fig. 8i**).

## Data availability

All data matrices representing mitral and tufted cell odor responses included in the analyses presented here are available upon request.

## Code availability

The code used for analysis is available upon request.

## Acknowledgements

The authors would like to acknowledge A.K. Dhawale, P. Gupta, G.B. Keller, P. Masset, M. Modi, S.D. Shea, P. Villar, A. Zhang, R.E. Egger, R.C. Muresan and members of the Albeanu lab for critical discussions, and M. Davis and R. Eifert for technical support. This work was supported by the following funding sources: a BBRF 2014 Young Investigator grant to H.C, a Junior Fellowship from the Simons foundation, NY to A.B and NSF IOS-1656830 and NIH R01DC014487-03 to D.F.A.

## Author contributions

H.C., A.B. and D.F.A. conceived the study and contributed to the design of experiments. H.C. performed the two-photon imaging experiments. H.C and A.B performed data analyses. A.B and D.F.A. wrote the manuscript.

## References

1. DiCarlo, J. J., Zoccolan, D. & Rust, N. C. How Does the Brain Solve Visual Object Recognition? Neuron 73, 415–434 (2012).

2. Rust, N. C. & DiCarlo, J. J. Selectivity and Tolerance (“Invariance”) Both Increase as Visual Information Propagates from Cortical Area V4 to IT. J. Neurosci. 30, 12978–12995 (2010).

3. Hong, H., Yamins, D. L. K., Majaj, N. J. & DiCarlo, J. J. Explicit information for category-orthogonal object properties increases along the ventral stream. Nat. Neurosci. 19, 613–622 (2016).

4. Yamins, D. L. K. et al. Performance-optimized hierarchical models predict neural responses in higher visual cortex. Proc. Natl. Acad. Sci. U. S. A. 111, 8619–8624 (2014).

5. Kar, K., Kubilius, J., Schmidt, K., Issa, E. B. & DiCarlo, J. J. Evidence that recurrent circuits are critical to the ventral stream’s execution of core object recognition behavior. Nat. Neurosci. 22, 974–983 (2019).

6. Bolding, K. A. & Franks, K. M. Complementary codes for odor identity and intensity in olfactory cortex. eLife https://elifesciences.org/articles/22630 (2017) doi:10.7554/eLife.22630.

7. Bolding, K. A. & Franks, K. M. Recurrent cortical circuits implement concentration-invariant odor coding. Science 361, (2018).

8. Stettler, D. D. & Axel, R. Representations of odor in the piriform cortex. Neuron 63, 854–864 (2009).

9. Pashkovski, S. L. et al. Structure and flexibility in cortical representations of odour space. Nature 583, 253–258 (2020).

10. Chae, H. et al. Mosaic representations of odors in the input and output layers of the mouse olfactory bulb. Nat. Neurosci. 22, 1306–1317 (2019).

11. Sirotin, Y. B., Shusterman, R. & Rinberg, D. Neural Coding of Perceived Odor Intensity,. eNeuro 2, (2015).

12. Moskowitz, H. R., Dravnieks, A. & Klarman, L. A. Odor intensity and pleasantness for a diverse set of odorants. Percept. Psychophys. 19, 122–128 (1976).

13. Mombaerts, P. Axonal wiring in the mouse olfactory system. Annu. Rev. Cell Dev. Biol. 22, 713–737 (2006).

14. Wilson, R. I. & Mainen, Z. F. Early events in olfactory processing. Annu. Rev. Neurosci. 29, 163–201 (2006).

15. Wiechert, M. T., Judkewitz, B., Riecke, H. & Friedrich, R. W. Mechanisms of pattern decorrelation by recurrent neuronal circuits. Nat. Neurosci. 13, 1003–1010 (2010).

16. Banerjee, A. et al. An Interglomerular Circuit Gates Glomerular Output and Implements Gain Control in the Mouse Olfactory Bulb. Neuron 87, 193–207 (2015).

17. Gschwend, O. et al. Neuronal pattern separation in the olfactory bulb improves odor discrimination learning. Nat. Neurosci. 18, 1474–1482 (2015).

18. Wanner, A. A. & Friedrich, R. W. Whitening of odor representations by the wiring diagram of the olfactory bulb. Nat. Neurosci. 23, 433–442 (2020).

19. Fukunaga, I., Berning, M., Kollo, M., Schmaltz, A. & Schaefer, A. T. Two distinct channels of olfactory bulb output. Neuron 75, 320–329 (2012).

20. Igarashi, K. M. et al. Parallel mitral and tufted cell pathways route distinct odor information to different targets in the olfactory cortex. J. Neurosci. Off. J. Soc. Neurosci. 32, 7970–7985 (2012).

21. Nagayama, S., Takahashi, Y. K., Yoshihara, Y. & Mori, K. Mitral and tufted cells differ in the decoding manner of odor maps in the rat olfactory bulb. J. Neurophysiol. 91, 2532–2540 (2004).

22. Jordan, R., Fukunaga, I., Kollo, M. & Schaefer, A. T. Active Sampling State Dynamically Enhances Olfactory Bulb Odor Representation. Neuron 98, 1214–1228.e5 (2018).

23. Kapoor, V., Provost, A. C., Agarwal, P. & Murthy, V. N. Activation of raphe nuclei triggers rapid and distinct effects on parallel olfactory bulb output channels. Nat. Neurosci. 19, 271–282 (2016).

24. Otazu, G. H., Chae, H., Davis, M. B. & Albeanu, D. F. Cortical Feedback Decorrelates Olfactory Bulb Output in Awake Mice. Neuron 86, 1461–1477 (2015).

25. Yamada, Y. et al. Context- and Output Layer-Dependent Long-Term Ensemble Plasticity in a Sensory Circuit. Neuron 93, 1198–1212.e5 (2017).

26. Burton, S. D. & Urban, N. N. Greater excitability and firing irregularity of tufted cells underlies distinct afferent-evoked activity of olfactory bulb mitral and tufted cells. J. Physiol. 592, 2097–2118 (2014).

27. Cavarretta, F. et al. Parallel odor processing by mitral and middle tufted cells in the olfactory bulb. Sci. Rep. 8, 7625 (2018).

28. Geramita, M. A., Burton, S. D. & Urban, N. N. Distinct lateral inhibitory circuits drive parallel processing of sensory information in the mammalian olfactory bulb. eLife 5, (2016).

29. Geramita, M. & Urban, N. N. Differences in Glomerular-Layer-Mediated Feedforward Inhibition onto Mitral and Tufted Cells Lead to Distinct Modes of Intensity Coding. J. Neurosci. Off. J. Soc. Neurosci. 37, 1428–1438 (2017).

30. Gire, D. H. et al. Mitral cells in the olfactory bulb are mainly excited through a multistep signaling path. J. Neurosci. Off. J. Soc. Neurosci. 32, 2964–2975 (2012).

31. Shepherd, G. M. Synaptic organization of the mammalian olfactory bulb. Physiol. Rev. 52, 864–917 (1972).

32. Nagayama, S. et al. Differential axonal projection of mitral and tufted cells in the mouse main olfactory system. Front. Neural Circuits 4, (2010).

33. M.D, G. M. S. The Synaptic Organization of the Brain. (Oxford University Press, USA, 2003).

34. Sosulski, D. L., Bloom, M. L., Cutforth, T., Axel, R. & Datta, S. R. Distinct representations of olfactory information in different cortical centres. Nature 472, 213–216 (2011).

35. Ghosh, S. et al. Sensory maps in the olfactory cortex defined by long-range viral tracing of single neurons. Nature 472, 217–220 (2011).

36. Boyd, A. M., Kato, H. K., Komiyama, T. & Isaacson, J. S. Broadcasting of cortical activity to the olfactory bulb. Cell Rep. 10, 1032–1039 (2015).

37. Markopoulos, F., Rokni, D., Gire, D. H. & Murthy, V. N. Functional properties of cortical feedback projections to the olfactory bulb. Neuron 76, 1175–1188 (2012).

38. Rothermel, M. & Wachowiak, M. Functional imaging of cortical feedback projections to the olfactory bulb. Front. Neural Circuits 8, 73 (2014).

39. Koulakov, A. A. & Rinberg, D. Sparse incomplete representations: a potential role of olfactory granule cells. Neuron 72, 124–136 (2011).

40. Grabska-Barwińska, A. et al. A probabilistic approach to demixing odors. Nat. Neurosci. 20, 98–106 (2017).

41. Gilbert, C. D. & Li, W. Top-down influences on visual processing. Nat. Rev. Neurosci. 14, 350–363 (2013).

42. Glickfeld, L. L., Andermann, M. L., Bonin, V. & Reid, R. C. Cortico-cortical projections in mouse visual cortex are functionally target specific. Nat. Neurosci. 16, 219–226 (2013).

43. Harris, K. D. & Mrsic-Flogel, T. D. Cortical connectivity and sensory coding. Nature 503, 51–58 (2013).

44. Petreanu, L. et al. Activity in motor-sensory projections reveals distributed coding in somatosensation. Nature 489, 299–303 (2012).

45. Marques, T., Nguyen, J., Fioreze, G. & Petreanu, L. The functional organization of cortical feedback inputs to primary visual cortex. Nat. Neurosci. 21, 757–764 (2018).

46. Leinweber, M., Ward, D. R., Sobczak, J. M., Attinger, A. & Keller, G. B. A Sensorimotor Circuit in Mouse Cortex for Visual Flow Predictions. Neuron 95, 1420–1432.e5 (2017).

47. Kwon, S. E., Yang, H., Minamisawa, G. & O’Connor, D. H. Sensory and decision-related activity propagate in a cortical feedback loop during touch perception. Nat. Neurosci. 19, 1243–1249 (2016).

48. Keller, A. J., Roth, M. M. & Scanziani, M. Feedback generates a second receptive field in neurons of the visual cortex. Nature 582, 545–549 (2020).

49. Kim, M.-H., Znamenskiy, P., Iacaruso, M. F. & Mrsic-Flogel, T. D. Segregated Subnetworks of Intracortical Projection Neurons in Primary Visual Cortex. Neuron 100, 1313–1321.e6 (2018).

50. Olsen, S. R., Bortone, D. S., Adesnik, H. & Scanziani, M. Gain control by layer six in cortical circuits of vision. Nature 483, 47–52 (2012).

51. Young, H., Belbut, B., Baeta, M. & Petreanu, L. Laminar-specific cortico-cortical loops in mouse visual cortex. bioRxiv 773085 (2019) doi:10.1101/773085.

52. Wilson, D. A. & Sullivan, R. M. Cortical processing of odor objects. Neuron 72, 506–519 (2011).

53. Choi, G. B. et al. Driving opposing behaviors with ensembles of piriform neurons. Cell 146, 1004–1015 (2011).

54. Miura, K., Mainen, Z. F. & Uchida, N. Odor representations in olfactory cortex: distributed rate coding and decorrelated population activity. Neuron 74, 1087–1098 (2012).

55. Roland, B., Deneux, T., Franks, K. M., Bathellier, B. & Fleischmann, A. Odor identity coding by distributed ensembles of neurons in the mouse olfactory cortex. eLife 6, (2017).

56. Babadi, B. & Sompolinsky, H. Sparseness and expansion in sensory representations. Neuron 83, 1213–1226 (2014).

57. Schaffer, E. S. et al. Odor Perception on the Two Sides of the Brain: Consistency Despite Randomness. Neuron 98, 736–742.e3 (2018).

58. Gottfried, J. A. Central mechanisms of odour object perception. Nat. Rev. Neurosci. 11, 628–641 (2010).

59. Kikuta, S. et al. Neurons in the anterior olfactory nucleus pars externa detect right or left localization of odor sources. Proc. Natl. Acad. Sci. 107, 12363–12368 (2010).

60. Esquivelzeta Rabell, J., Mutlu, K., Noutel, J., Martin del Olmo, P. & Haesler, S. Spontaneous Rapid Odor Source Localization Behavior Requires Interhemispheric Communication. Curr. Biol. 27, 1542–1548.e4 (2017).

61. Oettl, L.-L. et al. Oxytocin Enhances Social Recognition by Modulating Cortical Control of Early Olfactory Processing. Neuron 90, 609–621 (2016).

62. Wang, C. Y., Liu, Z., Ng, Y. H. & Südhof, T. C. A Synaptic Circuit Required for Acquisition but Not Recall of Social Transmission of Food Preference. Neuron 107, 144–157.e4 (2020).

63. Chen, T.-W. et al. Ultrasensitive fluorescent proteins for imaging neuronal activity. Nature 499, 295–300 (2013).

64. Meredith, M. Patterned response to odor in mammalian olfactory bulb: the influence of intensity. J. Neurophysiol. 56, 572–597 (1986).

65. Fusi, S., Miller, E. K. & Rigotti, M. Why neurons mix: high dimensionality for higher cognition. Curr. Opin. Neurobiol. 37, 66–74 (2016).

66. Barak, O., Rigotti, M. & Fusi, S. The sparseness of mixed selectivity neurons controls the generalization-discrimination trade-off. J. Neurosci. Off. J. Soc. Neurosci. 33, 3844–3856 (2013).

67. Kobak, D. et al. Demixed principal component analysis of neural population data. eLife 5, e10989 (2016).

68. Rao, R. P. & Ballard, D. H. Predictive coding in the visual cortex: a functional interpretation of some extra-classical receptive-field effects. Nat. Neurosci. 2, 79–87 (1999).

69. Olshausen, B. A. & Field, D. J. Sparse coding with an overcomplete basis set: a strategy employed by V1? Vision Res. 37, 3311–3325 (1997).

70. Rozell, C. J., Johnson, D. H., Baraniuk, R. G. & Olshausen, B. A. Sparse Coding via Thresholding and Local Competition in Neural Circuits. Neural Comput. 20, 2526–2563 (2008).

71. Boyd, A. M., Sturgill, J. F., Poo, C. & Isaacson, J. S. Cortical feedback control of olfactory bulb circuits. Neuron 76, 1161–1174 (2012).

72. Keller, G. B. & Mrsic-Flogel, T. D. Predictive Processing: A Canonical Cortical Computation. Neuron 100, 424–435 (2018).

73. Uchida, N., Kepecs, A. & Mainen, Z. F. Seeing at a glance, smelling in a whiff: rapid forms of perceptual decision making. Nat. Rev. Neurosci. 7, 485–491 (2006).

74. Andres, K. H. Anatomy and ultrastructure of the olfactory bulb in fish, amphibia, reptiles, birds, and mammals. in CIBA Foundation Symposium on Taste and Smell in Vertebrates 177–194 (Churchill).

75. Schwarz, D. et al. Architecture of a mammalian glomerular domain revealed by novel volume electroporation using nanoengineered microelectrodes. Nat. Commun. 9, 1–14 (2018).

76. Rokni, D., Hemmelder, V., Kapoor, V. & Murthy, V. N. An olfactory cocktail party: figure-ground segregation of odorants in rodents. Nat. Neurosci. 17, 1225–1232 (2014).

77. Haberly, L. B. Parallel-distributed processing in olfactory cortex: new insights from morphological and physiological analysis of neuronal circuitry. Chem. Senses 26, 551–576 (2001).

78. Mathis, A., Rokni, D., Kapoor, V., Bethge, M. & Murthy, V. N. Reading Out Olfactory Receptors: Feedforward Circuits Detect Odors in Mixtures without Demixing. Neuron 91, 1110–1123 (2016).

79. Uchida, N. & Mainen, Z. F. Speed and accuracy of olfactory discrimination in the rat. Nat Neurosci 6, 1224–1229 (2003).

80. Zariwala, H. A., Kepecs, A., Uchida, N., Hirokawa, J. & Mainen, Z. F. The limits of deliberation in a perceptual decision task. Neuron 78, 339–351 (2013).

81. Homma, R., Cohen, L. B., Kosmidis, E. K. & Youngentob, S. L. Perceptual stability during dramatic changes in olfactory bulb activation maps and dramatic declines in activation amplitudes. Eur. J. Neurosci. 29, 1027–1034 (2009).

82. Parthasarathy, K. & Bhalla, U. S. Laterality and Symmetry in Rat Olfactory Behavior and in Physiology of Olfactory Input. J. Neurosci. 33, 5750–5760 (2013).

83. Rajan, R., Clement, J. P. & Bhalla, U. S. Rats Smell in Stereo. Science 311, 666–670 (2006).

84. Feinberg, E. H. & Meister, M. Orientation columns in the mouse superior colliculus. Nature 519, 229–232 (2015).

85. May, P. J. The mammalian superior colliculus: laminar structure and connections. in Progress in Brain Research (ed. Büttner-Ennever, J. A.) vol. 151 321–378 (Elsevier, 2006).

86. Beltramo, R. & Scanziani, M. A collicular visual cortex: Neocortical space for an ancient midbrain visual structure. Science 363, 64–69 (2019).

87. Evans, D. A. et al. A synaptic threshold mechanism for computing escape decisions. Nature 558, 590–594 (2018).

88. Li, W. L. et al. Adult-born neurons facilitate olfactory bulb pattern separation during task engagement. eLife 7, (2018).

89. Lepousez, G. et al. Olfactory learning promotes input-specific synaptic plasticity in adult-born neurons. Proc. Natl. Acad. Sci. U. S. A. 111, 13984–13989 (2014).

90. Kay, L. M. & Laurent, G. Odor- and context-dependent modulation of mitral cell activity in behaving rats. Nat. Neurosci. 2, 1003–1009 (1999).

91. Frank, T., Mönig, N. R., Satou, C., Higashijima, S.-I. & Friedrich, R. W. Associative conditioning remaps odor representations and modifies inhibition in a higher olfactory brain area. Nat. Neurosci. 22, 1844–1856 (2019).

92. Hagiwara, A., Pal, S. K., Sato, T. F., Wienisch, M. & Murthy, V. N. Optophysiological analysis of associational circuits in the olfactory cortex. Front. Neural Circuits 6, (2012).

93. Mori, K., Kishi, K. & Ojima, H. Distribution of dendrites of mitral, displaced mitral, tufted, and granule cells in the rabbit olfactory bulb. J. Comp. Neurol. 219, 339–355 (1983).

94. Oswald, A.-M. & Urban, N. N. There and Back Again: The Corticobulbar Loop. Neuron 76, 1045–1047 (2012).

95. Brunjes, P. C., Illig, K. R. & Meyer, E. A. A field guide to the anterior olfactory nucleus (cortex). Brain Res. Brain Res. Rev. 50, 305–335 (2005).

96. Aqrabawi, A. J. & Kim, J. C. Hippocampal projections to the anterior olfactory nucleus differentially convey spatiotemporal information during episodic odour memory. Nat. Commun. 9, 2735 (2018).

97. Peron, S. P., Freeman, J., Iyer, V., Guo, C. & Svoboda, K. A Cellular Resolution Map of Barrel Cortex Activity during Tactile Behavior. Neuron 86, 783–799 (2015).

98. Kerlin, A. M., Andermann, M. L., Berezovskii, V. K. & Reid, R. C. Broadly Tuned Response Properties of Diverse Inhibitory Neuron Subtypes in Mouse Visual Cortex. Neuron 67, 858–871 (2010).

99. Khan, A. G. et al. Distinct learning-induced changes in stimulus selectivity and interactions of GABAergic interneuron classes in visual cortex. Nat. Neurosci. 21, 851–859 (2018).

100. Hopfield, J. J. Pattern recognition computation using action potential timing for stimulus representation. Nature 376, 33–36 (1995).

101. Tank, D. W. & Hopfield, J. J. Neural computation by concentrating information in time. Proc. Natl. Acad. Sci. U. S. A. 84, 1896–1900 (1987).

102. Gollisch, T. & Meister, M. Rapid neural coding in the retina with relative spike latencies. Science 319, 1108–1111 (2008).

103. Wilson, C. D., Serrano, G. O., Koulakov, A. A. & Rinberg, D. A primacy code for odor identity. Nat. Commun. 8, 1477 (2017).

