## Extended Data Materials for "A non-canonical feedforward pathway for computing odor identity"

10 Extended Data Figures
1 Extended Data Table
References

### Extended Data Figure 1

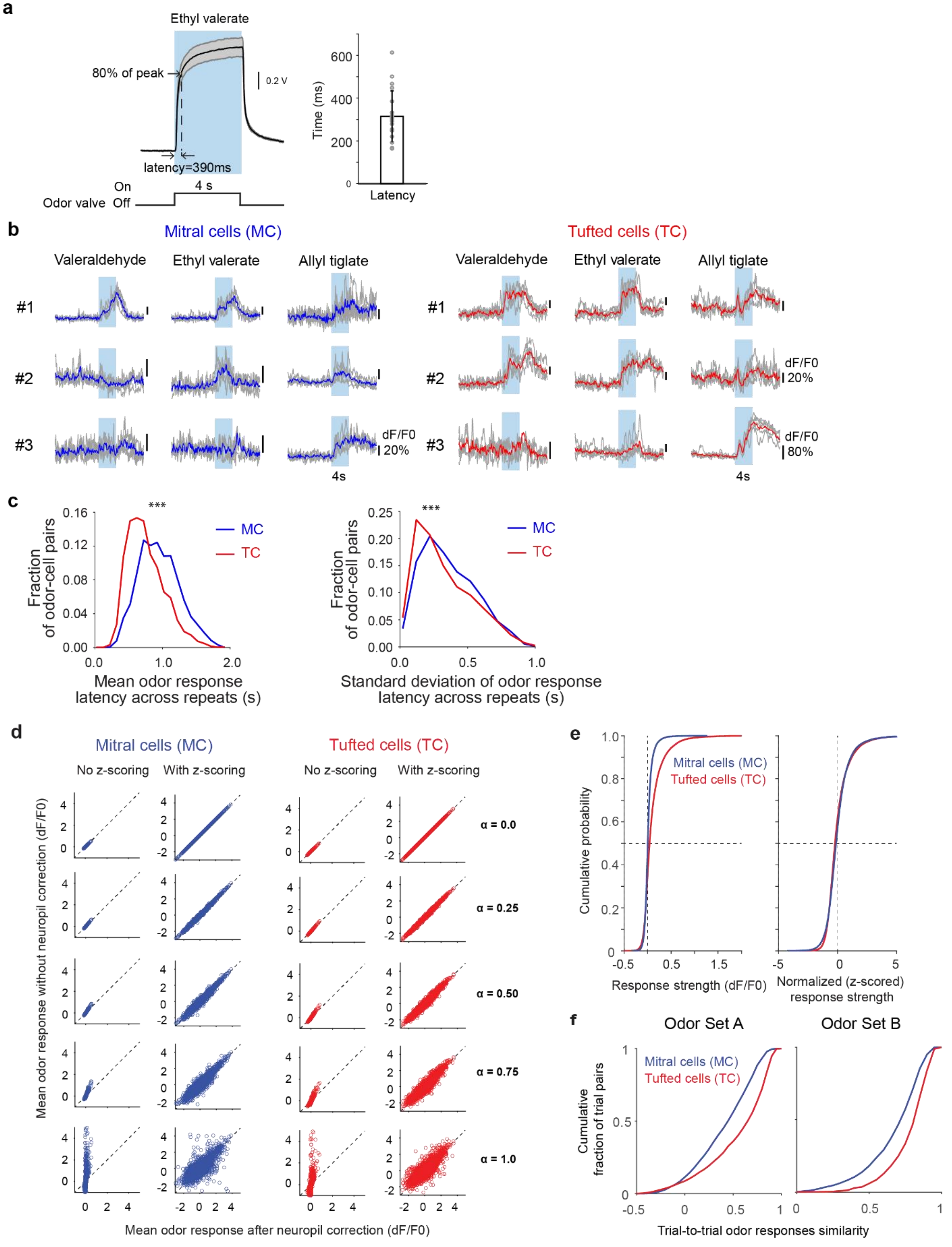

**Extended Data Figure 1. Calibration of odor delivery, latency, reproducibility of mitral and tufted cells odor responses in awake naïve mice, and normalization of mitral and tufted cells odor response strength and variability of responses across trials**

**a.** (*Left*) Photo-ionization device (PID) average trace (black) and standard deviation (gray, 5 repeats) for an example odor in the panel (ethyl valerate, $10^{-3}$ oil dilution). Blue color indicates duration of stimulus delivery (odor valve ON, 4s). Latency is calculated as the interval from odor valve opening command until signal reaches 80% of its peak value. (*Right*) Average latency (315±119ms standard deviation, STD) of 5 odors across 4 dilutions (Odor Set A).

**b.** Single trial responses obtained via multiphoton imaging of GCaMP6f signals of three example mitral (*Left*) and tufted (*Right*) cells to three odors (valeraldehyde, ethyl valerate and allyl tiglate, $10^{-2}$ oil dilution); shaded area marks duration of odor presentation (4s), blue/red lines mark the average change in fluorescence ($dF/F_0$) across four trials; gray traces correspond to individual trials.

**c.** (*Left*) Distribution of mean response latency across odor responses in mitral versus tufted cells (MC: Avg. 949.4±6.3ms standard error of the mean, SEM, 2,069 odor-cell pairs; TC: Avg. 754.9±4.1ms, 4,460 odor-cell pairs, $p<0.001$, Wilcoxon rank sum test). (*Right*) Standard Deviation of response latency of cell-odor response pairs across repeats (MC: Avg. STD 357.6ms; TC: Avg. STD 332.6ms, 2,069 odor-cell pairs, $p<0.001$, Wilcoxon rank sum test); blue traces indicate mitral cells and red traces tufted cells.

**d.** (*Left*) Average fluorescence responses ($dF/F_0$) without versus with- neuropil correction (Methods, $F_{ROI\text{-}corrected} = F_{ROI} - \alpha F_{neuropil}$) in a non-z-scored and z-scored example mitral cell field of view, while varying the α parameter between 0 to 1 in 0.25 increments; each dot corresponds to one odor-cell pair; dashed line marks unity slope. (*Right*) same for tufted cells.

**e.** Cumulative histograms of raw (*Left*) and z-scored normalized (*Right*) response strength for the mitral and tufted cells sampled (n = 447 MC, 558 TC, 5 odorants across 4 different concentrations); vertical dashed lines mark z-score=0; horizontal dashed lines mark 0.5 cumulative probability.

**f.** Trial-to-trial odor response (z-scored) similarity for the concentration data-set (*Left, Odor Set A*) and large odor data-set (*Right, Odor Set B*) for both mitral (*blue*) and tufted (*red*) cells. Cumulative histograms aggregated across all stimuli and fields-of-view; $p<0.001$, Wilcoxon rank-sum test.

#### Extended Data Figure 2

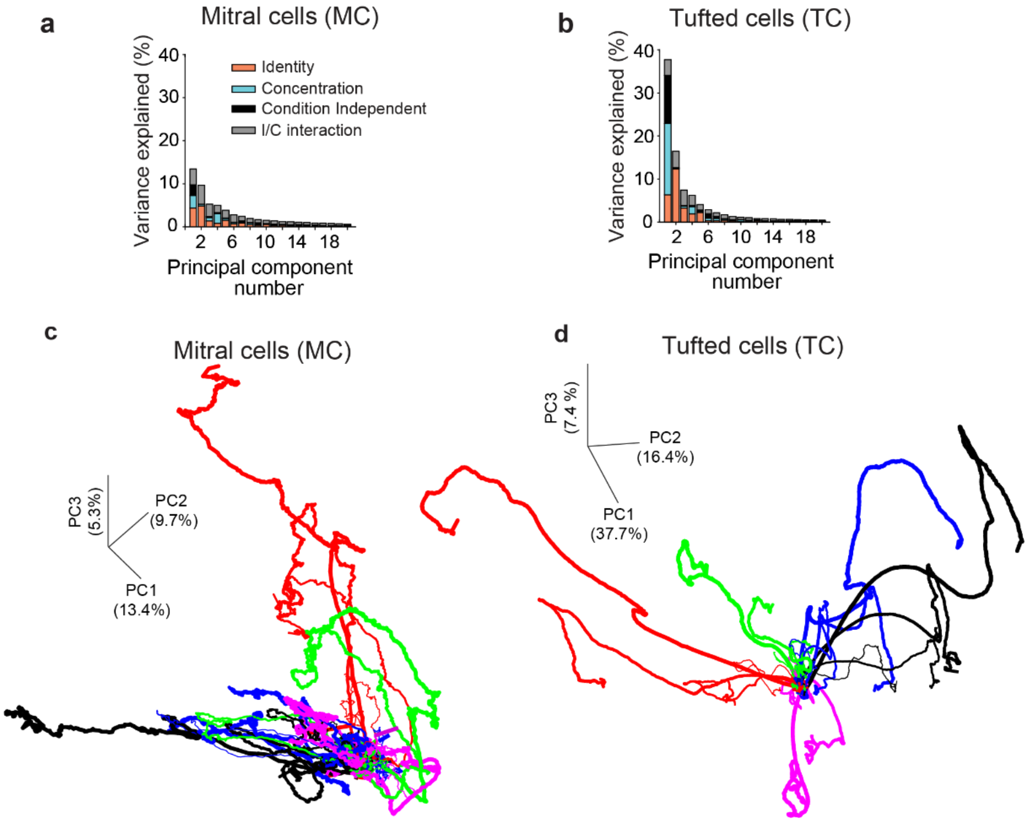


**Extended Data Figure 2. Principal component analysis (PCA) on mitral and tufted cell ensemble odor responses across different concentrations**

**a-b.** For mitral (**a**) and tufted (**b**) cell ensembles, variance explained by the top 20 principal components identified using PCA decomposed into four categories: 'odor identity', 'odor concentration', 'interaction between identity and concentration' (I/C interaction) and 'condition independent'; n = 447 MCs and 458 TCs; stimuli: 5 odors, 4 concentrations.

**c.** Mitral cell ensemble trajectories in the neural state space defined by the top three principal components in descending order of total variance explained. Different colors denote different odorants while increasing thickness indicates increasing concentration.

**d.** Same as **c**, except for tufted cell ensembles.

#### Extended Data Figure 3

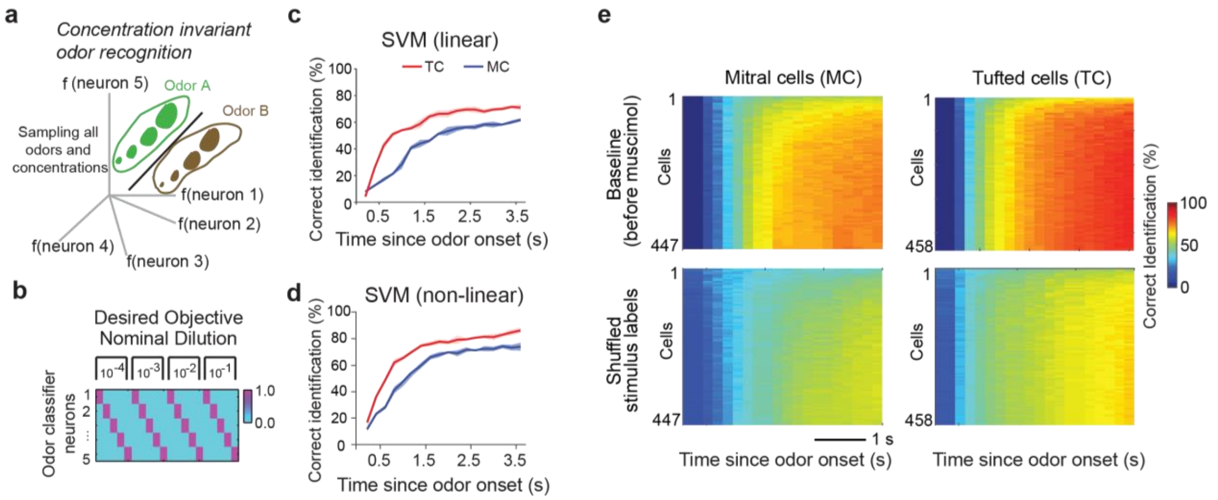


**Extended Data Figure 3. Concentration invariant odor decoding performance depends on concentration representations**

**a.** *Concentration-invariant odor recognition*. In this decoding scheme, each stimulus occupies a distinct region of the neural state space and all concentrations of a given odorant need to be grouped together as one odor object.

***b.*** Set-up of the decoding strategy where hypothetical classifier neurons (one for each odorant) signal the presence (value =1) of their corresponding odorant for all four sampled concentrations, and its absence (value = 0) for all other odorants in the panel.

**c.**,**d.** Cross-validated classification performance in the *concentration-invariant odor* recognition decoding scheme using a linear support vector machine (SVM) decoder (**c**, see Methods) and a non-linear SVM decoder (**d**, see Methods) averaged across all five odorants for mitral (*blue*) and tufted (*red*) cells.

**e**. 2D classification performance map for mitral cells (MC, *Top Left*) and tufted cells (TC, *Top Right*) with increasing cell number (ordinate) and time elapsed (abscissa) using a non-linear SVM decoder. (*Bottom*) Same as *Top*, except concentration labels of each odorant were shuffled independently between the training and the testing conditions, thereby challenging the decoder to learn arbitrary grouping of 4 random stimuli at a time. As expected, the shuffled classifier performance was significantly lower, highlighting that classification accuracy of invariant odor identification depends on the learned concentration labels.

**Extended Data Figure 4**

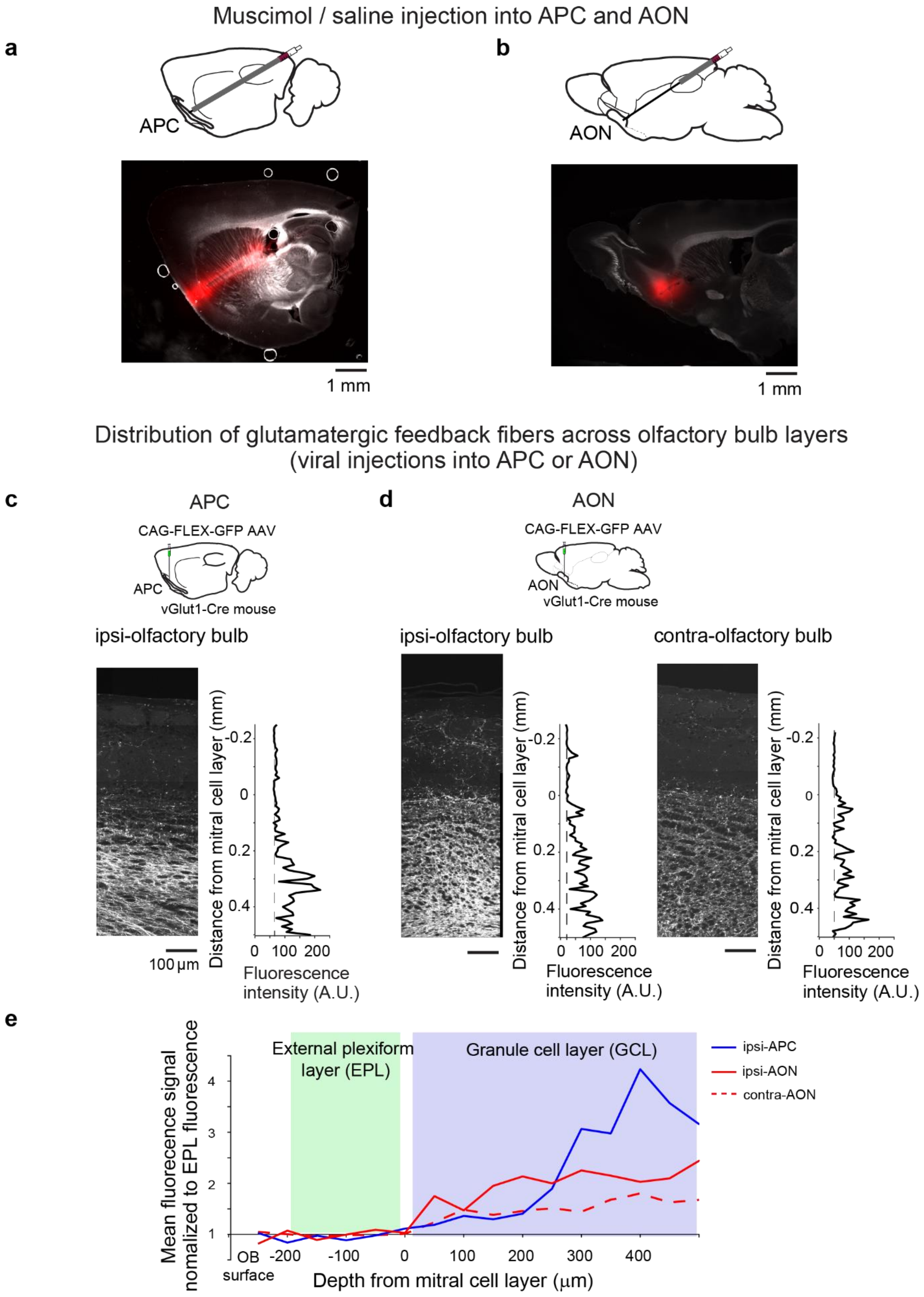

**Extended Data Figure 4. Histology of muscimol injections in the olfactory cortex (anterior piriform cortex and anterior olfactory nucleus) and the distribution of glutamatergic cortical feedback fibers to the olfactory bulb**

**a**., **b**. Sites of muscimol/saline injection in the anterior piriform cortex (APC, **a**) and anterior olfactory nucleus (AON), posterior part, AOP (**b**) in TBET-Cre x AI95 (GCaMP6f) mice. (*Top*) Cartoon sagittal views of the brain depicting the locations of the guide cannula. (*Bottom*) Example brain slices with the cannula track and injection sites outlined by fluorescent muscimol. Fluorescent muscimol also marks the cannula tract; staining was obtained by gently retrieving the cannula while injecting.

**c**., **d**. Distribution of glutamatergic cortical feedback fibers across the olfactory bulb layers. For illustration purposes, cortical-bulbar feedback was labeled in vGlut1-Cre mice by injection of CAG-FLEX-GFP AAV in the ipsi-APC (**c**) as well as the ipsi- or contra-AON (**d**). vGlut-cre mice were used here to minimize viral expression in olfactory bulb somata due to infection of migrating adult-born interneurons. Previous work indicates that cortical-bulbar feedback is mostly glutamatergic[1–7].

**e.** Distribution of mean fluorescence signal of cortical feedback fibers in different layers of the olfactory bulb normalized to their average fluorescence in the external plexiform layer (EPL); zero marks the mitral cell layer; normalized fluorescence is plotted for feedback fibers from ipsi-APC, ipsi-AON and contra-AON.

### Extended Data Figure 5

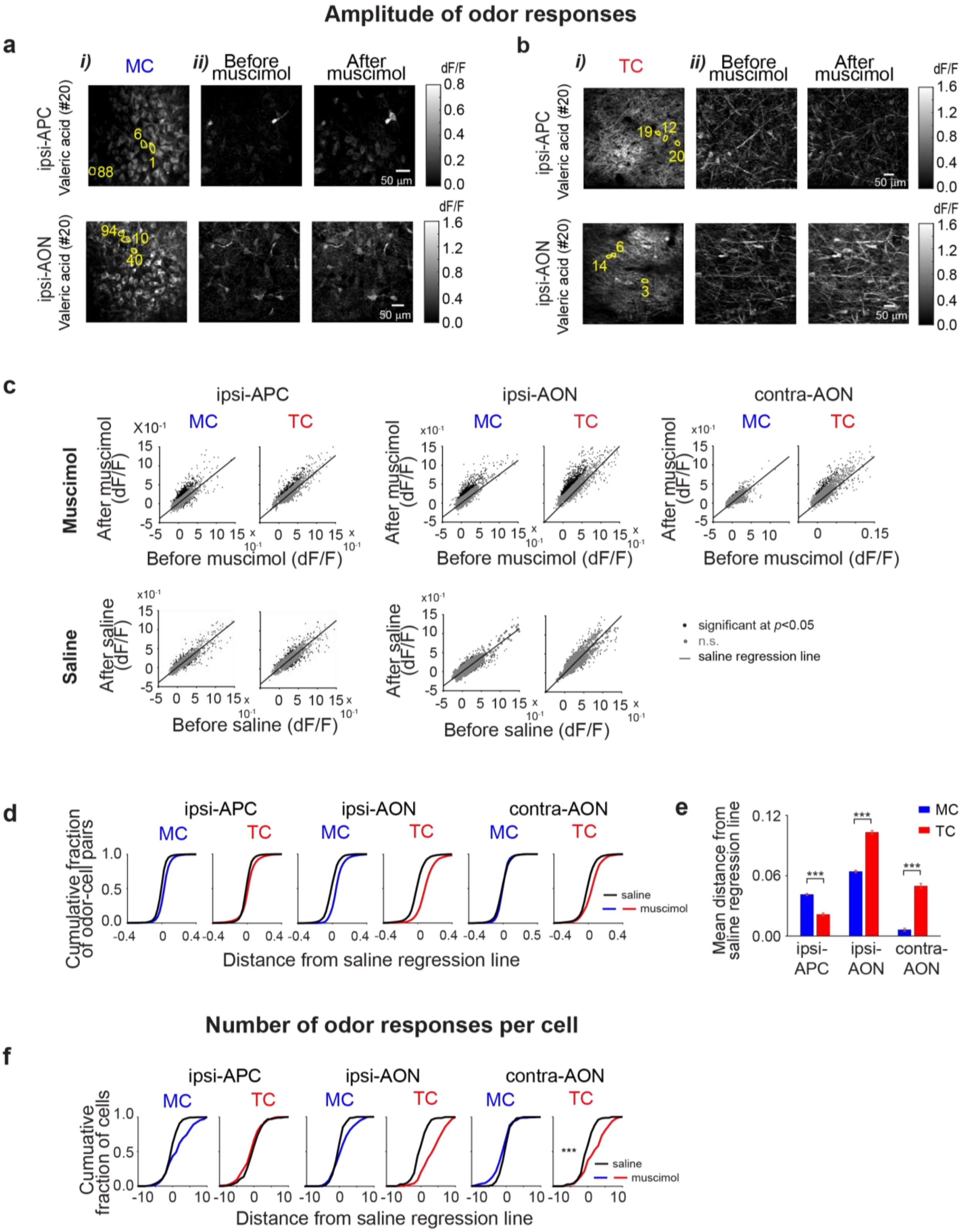

**Extended Data Figure 5. Example odor responses and quantification of differential effects of cortical feedback suppression on mitral and tufted cells**

**a.**,**b**. *i*) Average resting fluorescence of example field of view containing mitral cells (MC) somata (~225µm from surface) and tufted cell (TC) somata in the external plexiform layer (~150µm from surface).

**a.**,**b**. *ii*) Ratio image showing average fluorescence change ($dF/F_0$) in response to valeric acid in the field of view shown in **a.**,**b**. *i)* before (*Left*) and after (*Right*) muscimol injection into APC (*Top*) or AON (*Bottom*).

**c**. Scatter plots showing the amplitude of mitral and tufted cell odor responses before versus after muscimol (*Top*) or saline (*Bottom*) injection into ipsi-APC (*Left*), ipsi-AON (*Center*) and contra-AON (*Right*); a saline regression line was obtained by minimizing the Euclidian distances to the cell-odor pairs included in the analysis. A 95% percentile confidence interval with respect to the saline regression line was imposed for calculating significance (black vs. gray dots) in muscimol injection experiments.

**d.** Cumulative plots of distance from saline regression for changes in mitral and tufted cell odor response amplitude ($dF/F_0$) after muscimol (blue: MC, red: TC) versus saline (black) injection into ipsi-APC (*Left*), ipsi-AON, (*Center*) or contra-AON, (*Right*).

**e**. *ii*) Summary of mean distance from saline regression line for changes in odor response amplitude ($dF/F_0$) of mitral (blue) and tufted cell (red) representations post muscimol injection into ipsi-APC (MC: muscimol 0.041±0.001, n=4,682 odor-cell pairs; TC: muscimol 0.022±0, n=4,316 odor-cell pairs; MC vs. TC), ipsi-AON (MC: muscimol 0.064±0.001, n=3,971 odor-cell pairs; TC: muscimol 0.103±0.002, n=3,777 odor-cell pairs; MC vs. TC), or contra-AON (MC: muscimol 0.006±0.002, n=1,656 odor-cell pairs; TC: muscimol 0.050±0.003, n=1,861 odor-cell pairs; MC vs. TC); *** marks $p<0.001$, Wilcoxon rank sum test. All numbers represent mean and SEM.

**f.** Cumulative plots of distance from saline regression for changes in the number of odor responses per cell after muscimol (blue: MC, red: TC) versus saline (black) injection into ipsi-APC (*Left*), ipsi-AON (*Center*) or contra-AON, (*Right*).

**Extended Data Figure 6**

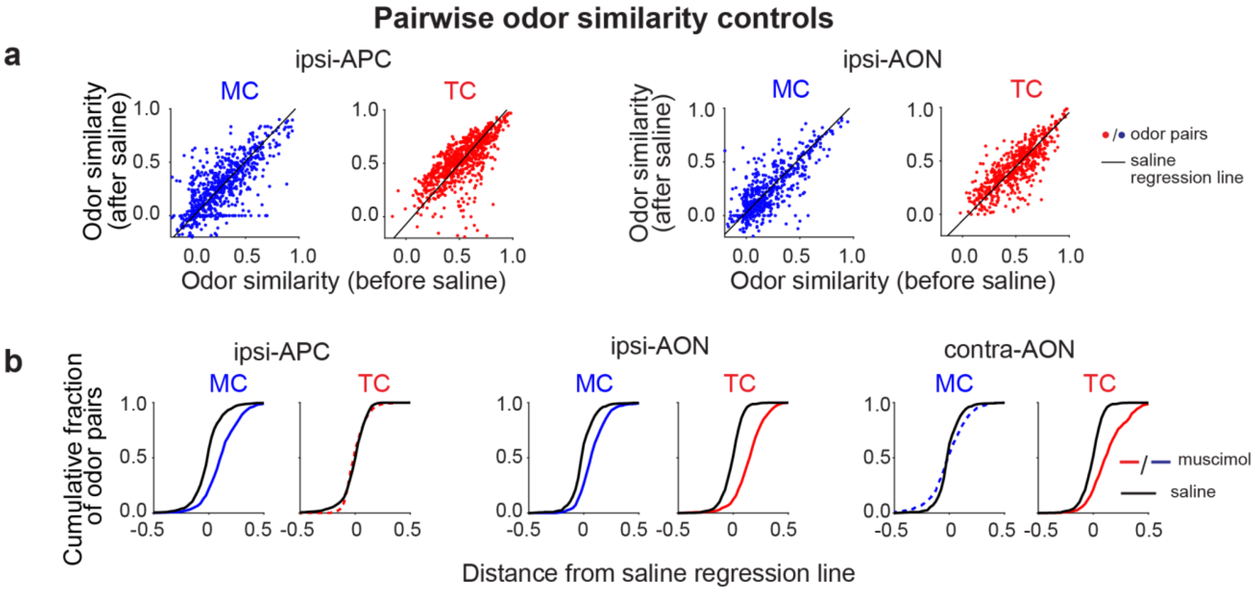


**Extended Data Figure 6. Saline controls for pairwise odor similarity representations in mitral and tufted cell ensembles before and after suppression of cortical bulbar feedback from the anterior piriform cortex or anterior olfactory nucleus**

**a.** Scatter plots of pairwise odor similarity of mitral (blue) and tufted cell (red) responses before and after saline injection into ipsi-APC (*Left*) and ipsi-AON (*Right*); each dot represents one odor-to-odor comparison before and after saline injection; combined responses from mitral or tufted cells across all sampled fields of view; ipsi-APC, MC: n=950 odor pairs from 5 FOVs; TC: n=950 odor pairs from 5 FOVs; ipsi-AON, MC: n=760 odor pairs from 4 FOVs; TC: n=760 odor pairs from 4 FOVs.

**b.** Cumulative distributions of distance with respect to saline regression line for mitral and tufted cell pairwise odor similarity after saline (black) and muscimol (blue: MC, red: TC) injection into ipsi-APC(*Left)*, ipsi-AON (*Center*) and contra-APC (*Right*). Dashed line denotes lack of significant difference with respect to the saline control.

### Extended Data Figure 7

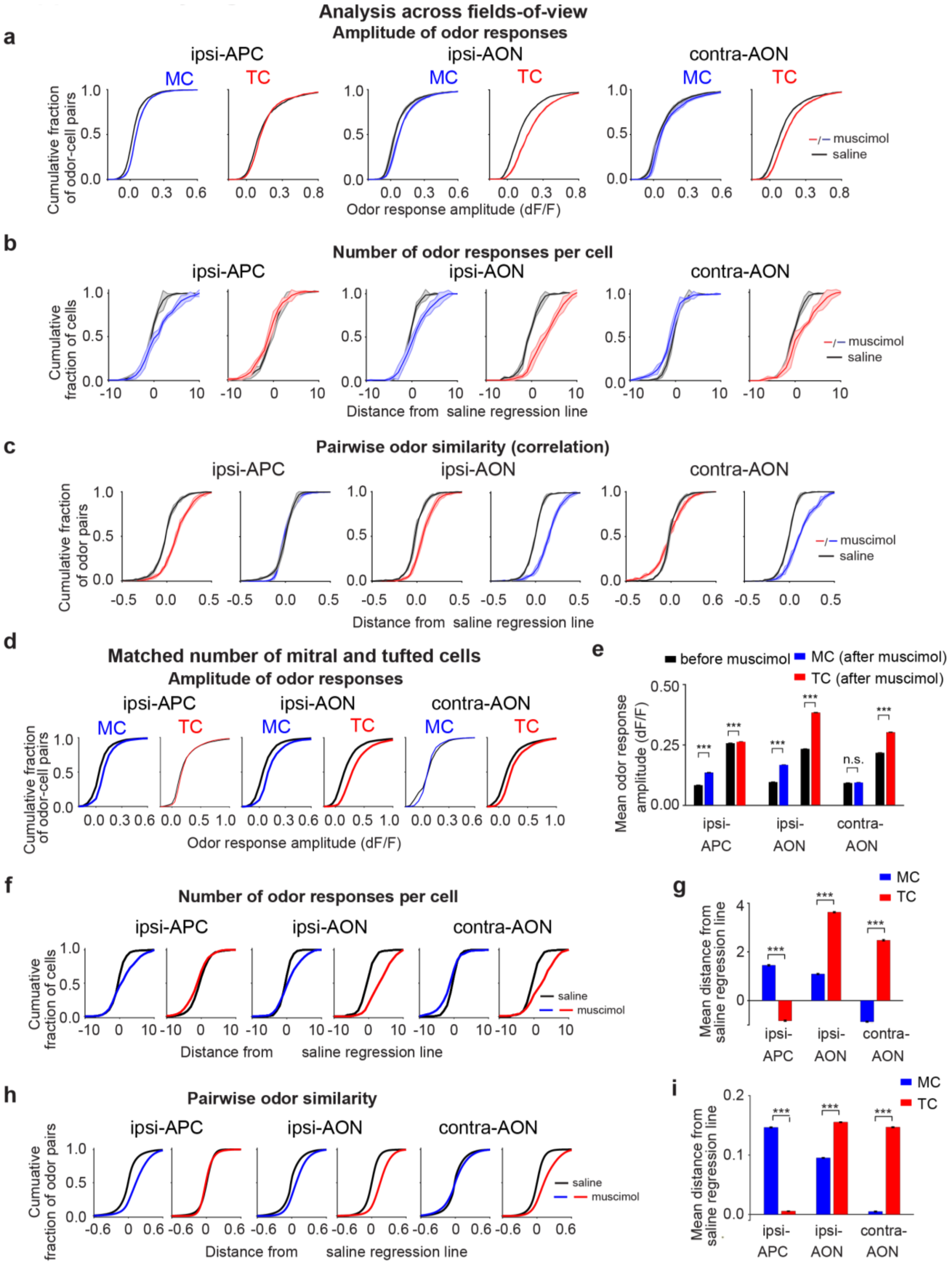

**Extended Data Figure 7. Odor responsiveness and pairwise odor similarity in *individual fields of view* and *matched number* bootstrap analyses of mitral and tufted cell representations before versus after suppression of cortical bulbar feedback from APC or AON**

**a.** Cumulative distribution of mitral or tufted cell-odor response pairs as function of $dF/F_0$ responses amplitude before (black) and after (blue: MC, red: TC) muscimol injection into ipsi-APC (*Left,* MC: n=5 FOVs; TC: n=5 FOVs), ipsi-AON (*Center,* MC: n=4 FOVs; TC: n=4 FOVs) and contra-AON (*Right,* MC: n=3 FOVs; TC: n=3 FOVs). Shaded area marks standard error across individual field of views.

**b.** Cumulative plots of distance from saline regression for changes in the number of odor responses per cell after muscimol (blue: MC, red: TC) versus saline (black) injection into ipsi-APC(*Left,* MC: n=5 FOVs; TC: n=5 FOVs), ipsi-AON (*Center,* MC: n=4 FOVs; TC: n=4 FOVs) and contra-AON (*Right,* MC: n=3 FOVs; TC: n=3 FOVs). Shaded area marks standard error across individual field of views.

**c.** Cumulative plots of distance distributions from saline regression of mitral (blue) and tufted cell (red) odor similarity distributions after muscimol versus saline injection into ipsi-APC (*Left,* MC: n=5 FOVs; TC: n=5 FOVs), ipsi-AON (*Center,* MC: n=4 FOVs; TC: n=4 FOVs) and contra-AON (*Right,* MC: n=3 FOVs, TC: n=3 FOVs). Shaded area marks standard error across individual field of views.

**d.** For each field of view, 40 mitral cells and 40 tufted cells were randomly selected and a bootstrap analysis ran for 100 iterations; for each iteration 4 FOVs were selected. Cumulative distribution of MC or TC-odor response pairs as function of $dF/F_0$ responses amplitude before (black) and after (blue: MC, red: TC) muscimol injection into ipsi-APC (*Left*), ipsi-AON (*Center*) and contra-AON (*Right*).

**e.** Summary of mean odor responses amplitude ($dF/F_0$) of mitral and tufted cells before (black) and after (blue: MC, red: TC) muscimol injection into ipsi-APC (MC: before 0.083±0.000, after 0.136±0. 000; TC: before 0.257±0. 000, after 0.263±0.039), ipsi-AON (MC: before 0.096±0.000, after 0.167±0.000; TC: before 0.233±0.037, after 0.383±0.000) and contra-AON (MC: before 0.092±0.000, after 0.094±0.000; TC: before 0.217±0.000, after 0.302±0.000). *** marks $p<0.001$, One-sided Wilcoxon rank sum test. All numbers represent mean and SEM.

**f.** Cumulative plots of distance from saline regression for changes in the number of odor responses per cell after muscimol (blue: MC, red: TC) versus saline (black) injection into ipsi-APC (*Left*), ipsi-AON (*Center*) and contra-AON (*Right*).

**g.** Summary of mean distance from saline regression line for changes in the number of odor responses per cell after muscimol (blue: MC, red: TC) into ipsi-APC (MC: muscimol 1.451±0.029; TC: muscimol -0.827±0.024; MC vs. TC: $p<0.001$), ipsi-AON (MC: muscimol 1.068±0.026; TC: muscimol 3.676±0.026; MC vs. TC: $p<0.001$) and contra-AON (MC: muscimol -0.846±0.023; TC: muscimol 2.480±0.029; MC vs. TC: $p<0.001$); *** marks $p<0.001$, Wilcoxon rank sum test. All numbers represent mean and SEM.

**h.** Cumulative plots of distance distributions from saline regression of mitral (blue) and tufted cell (red) odor similarity distributions after muscimol injection into ipsi-APC (*Left*), ipsi-AON (*Center*) and contra-AON (*Right*).

**i.** Summary of mean distance from saline regression line for pairwise odor similarity of mitral (blue) and tufted cell (red) representations before and after muscimol injection into ipsi.-APC (MC: 0.147±0.000, TC: 0.006±0.000; MC vs. TC: $p<0.001$), ipsi-AON (MC: 0.095±0.000, TC: 0.155±0.000; MC vs. TC: $p<0.001$) and contra-AON (MC: 0.005±0.000, TC: 0.147±0.000; MC vs. TC: $p<0.001$); *** marks $p<0.001$, Wilcoxon rank sum test. All numbers represent mean and SEM.

### Extended Data Figure 8

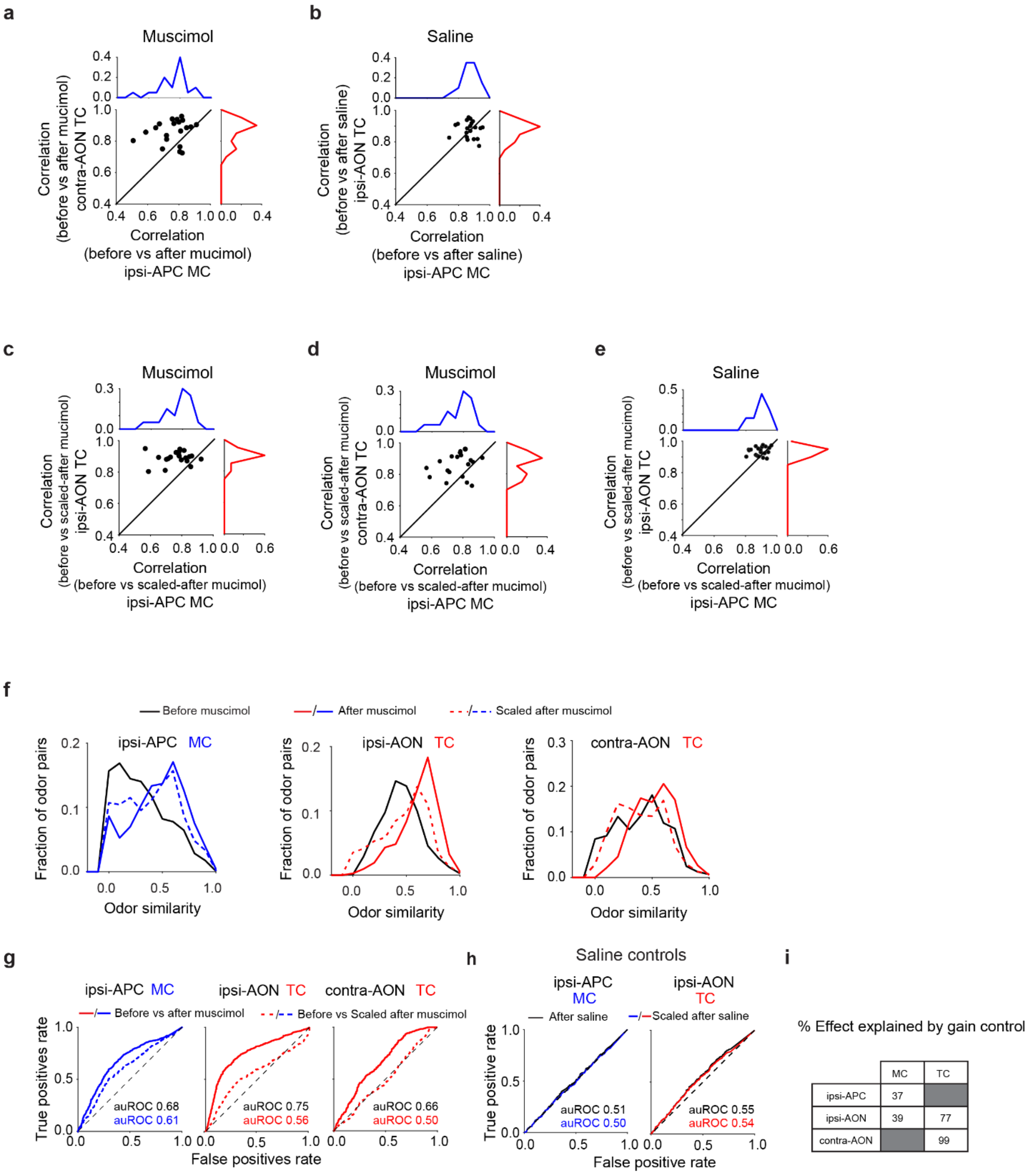

**Extended Data Figure 8. Cortical feedback controls the gain of tufted cell odor representations, and restructures mitral cell responses beyond simple scaling**

**a-b.** Scatter plot and histograms for pairwise similarity of mitral (*blue*) and tufted (*red*) cell response spectra to same odor before versus after muscimol (**a**) injection into ipsilateral APC and contralateral AON or saline (**b**) injection into ipsilateral APC and ipsilateral AON.

**c-e.** Scatter plot and histograms for pairwise similarity of mitral and tufted cell response spectra to same odor before versus scaled down after muscimol injection into ipsilateral APC and ipsilateral AON (**c**), scaled down after muscimol injection into ipsilateral APC and contralateral AON (**d**), or scaled after saline injection into ipsilateral APC and ipsilateral AON (**e**).

**f.** Distributions of pairwise odor similarity in mitral cells and tufted cells representations before (*black*) and after suppression of ipsilateral APC (*blue*, *mitral cells*), ipsilateral and contralateral AON (*red*, *tufted cells*) activity by muscimol injection (*solid lines*), as well as after scaling down post-muscimol responses (*dashed lines*) so as to match the mean of before-muscimol odor responses.

**g.** Receiver Operating Characteristic, ROC analysis for mitral cells (*blue*) and tufted cells (*red*) comparing the separability of *after muscimol* versus *before-muscimol* mitral cell pairwise odor similarity distributions (*solid lines*) and *scaled down after-muscimol* versus *before-muscimol* odor similarity distributions (*dashed lines*). auROC – area under the ROC curve. Consistent with a predominantly gain control role of AON feedback onto tufted cells, 'down-scaled' after- tufted cell response distributions were only marginally different from, or overlapped with, the pre-inactivation distributions (ipsi- AON auROC = 0.56, contra- AON auROC = 0.50). In contrast, for mitral cells, scaling down responses could not readily account for the observed differences before- and after- APC suppression (ipsi- APC auROC = 0.61)

**h.** (*Left*) ROC analysis comparing the separability of *after-saline* injection versus *before-saline* mitral cell pairwise odor similarity distributions (black, auROC =0.50) and *scaled down after-saline* versus *before-saline* mitral cell odor similarity distributions (blue, auROC=0.51). (*Right*) ROC analysis comparing the separability of *after-saline* injection versus *before-saline* tufted cell pairwise odor similarity distributions (black, auROC =0.55) and *scaled down after-saline* versus *before-saline* tufted cell odor similarity distributions (red, auROC=0.54). As expected, in saline injection controls the three distributions were indistinguishable for both mitral and tufted cells.

**i.** Percentage change in auROC which can be explained by scaling down *after-muscimol* odor responses for mitral and tufted cell odor similarity distributions respectively, when suppressing ipsi-APC, ipsi-AON and contra-AON. Since ipsi-APC suppression did not significantly alter tufted cell odor representations, and suppression of contra-AON did not impact MC representations, these table entries were left blank. On average, a substantially larger fraction of cortical feedback action on tufted versus mitral cell ensembles could be accounted for by gain control scaling (>80% vs. ~40%).

**Extended Data Figure 9**

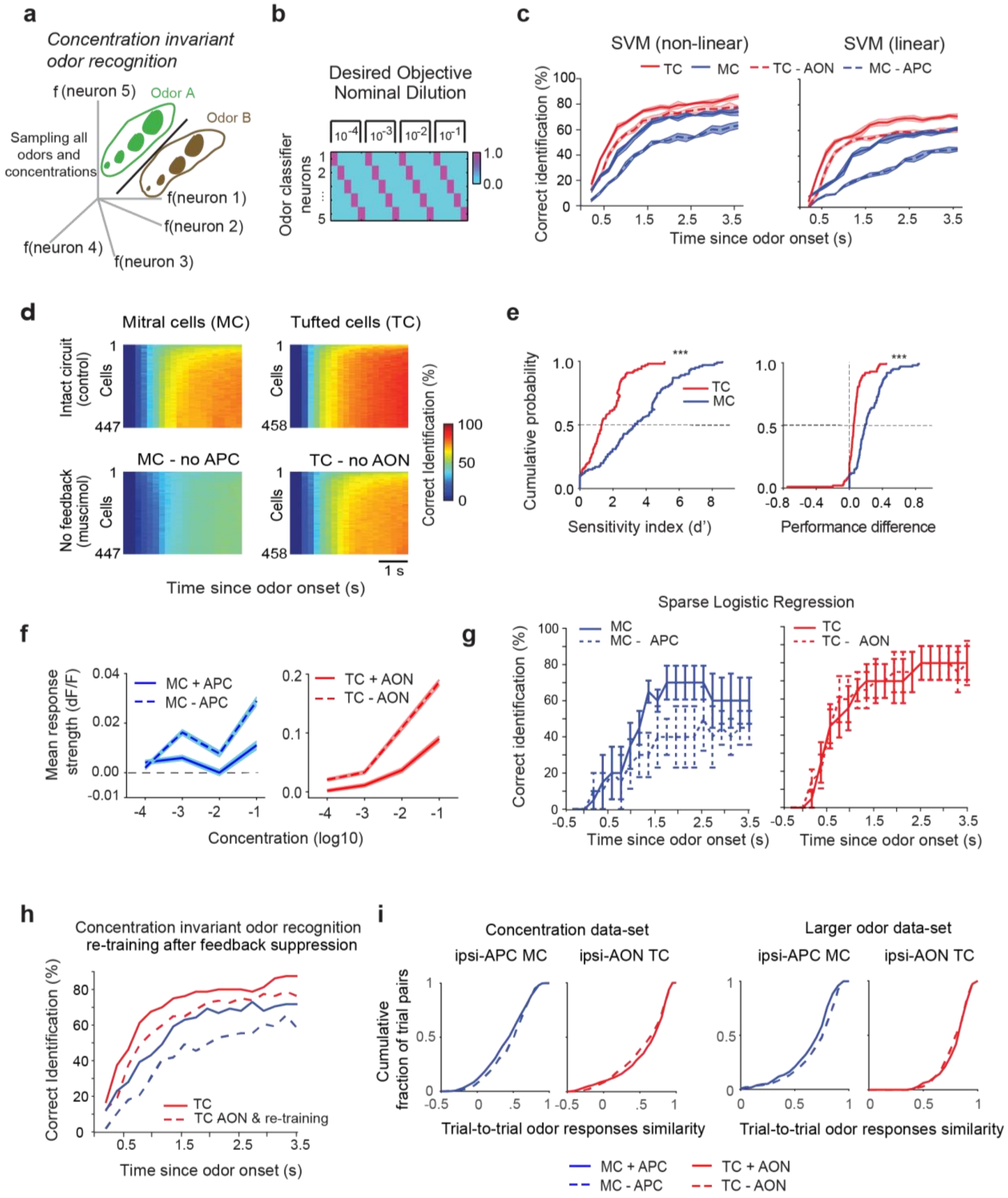

**Extended Data Figure 9. Effect of cortical feedback on concentration invariant odor recognition**

**a.** Cartoon schematics of *concentration-invariant odor recognition* decoding. Each stimulus occupies a distinct region of the neural state space and all concentrations of a given odorant need to be grouped together by the classifier.

**b.** The decoding objective function where one hypothetical classifier neuron signals the presence (value =1) of its corresponding odorant for each of four concentrations sampled and absence (value = 0) when any other odor in the panel is delivered instead. Cross-validated performance is tested on held-out trials previously not used to train.

**c.** Cross-validated classification performance of a non-linear polynomial kernel for the support vector machine (SVM, *Top*, Methods) and a linear SVM (*Bottom*, Methods) as a function of time for mitral cells (*blue*) and tufted cells (*red*) with (*solid line*) and without feedback (*dashed line*) from preferred cortical targets (APC for mitral cells and AON for tufted cells).

**d.** 2-D classification performance color map for all four experimental conditions as a function of time (abscissa, bin size = 200 ms), while varying the number of neurons included in the analysis using bootstrap re-sampling (ordinate, bin size=5 neurons).

**e.** For both mitral (*blue*) and tufted (*red*) cells, classifier performance difference with and without cortical feedback is quantified using d-prime or a performance difference index (Methods). In both cases, the performance drop after cortical feedback suppression is significantly higher for mitral cells than tufted cells. *** indicates $p<0.001$, paired t-test.

**f.** Population concentration response averaged across all mitral (*Left*, n = 447, blue) cells and tufted cells (*Right*, n = 458 cells) in the presence (solid line) or after suppression (dashed line) of cortical feedback. Concentration is represented using a log scale.

**g.** Cross-validated classification performance using a sparse logistic regression (SLR) decoder (Methods) in the *concentration-invariant odor recognition* decoding scheme for mitral cell (*Left, solid blue line*) and tufted cell (*Right*, *solid red line*) ensemble in the presence and after suppression of cortical feedback (*dashed lines*) from APC and AON respectively.

**h.** Cross-validated classification performance in *concentration-invariant odor recognition* for mitral (*red*) and tufted (*blue*) cells using a non-linear SVM decoder. To assess the decoding accuracy after suppression of APC and AON cortical feedback respectively (*dashed lines*), the tufted and mitral cell based decoders were re-trained separately with identical parameters.

**i.** Trial-to-trial odor response (z-scored) similarity for the concentration data-set (*Left, Odor Set A*) and large odor data-set (*Right, Odor Set B*) for both mitral (*Blue*) and tufted (*Red*) cells before (*solid lines*) and after cortical inactivation (*dashed lines*). Cumulative histograms aggregated across all stimuli and fields-of-view.

**Extended Data Figure 10**

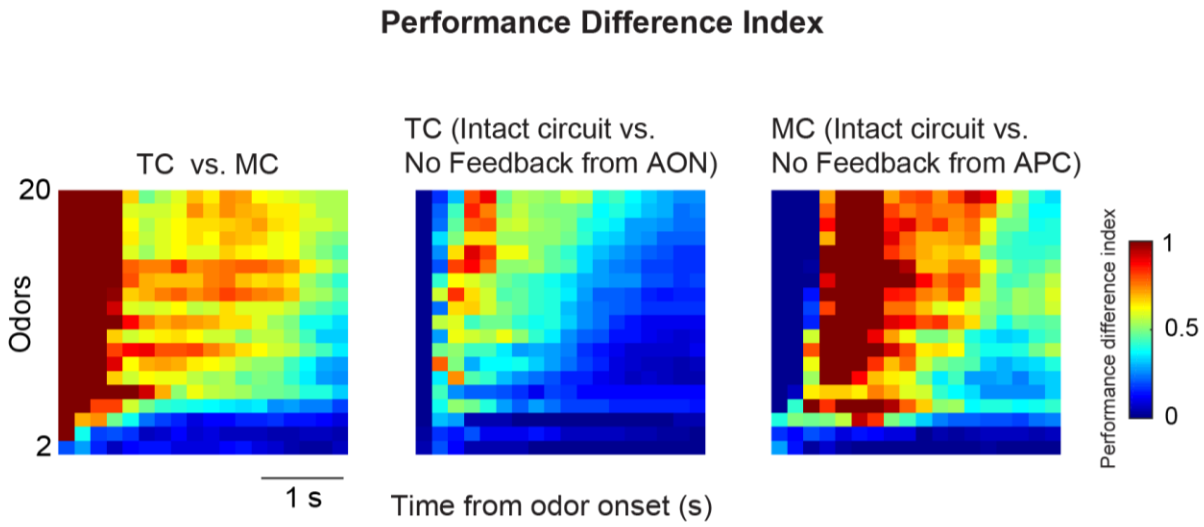


**Extended Data Figure 10. Effect of cortical feedback suppression on mitral and tufted cell ensemble decoding in discrimination across a larger odor panel**

2D performance difference index map with increasing number of odor distractors (ordinate) and increasing time from odor onset (abscissa). (*Left*) Tufted cells (TC) vs. mitral cells (MC) based decoders; positive values indicate tufted cells performance is higher than mitral cells. (*Center*) Change in performance of tufted cells decoders before vs. after suppression of AON feedback; (*Right*) Change in performance of mitral cells decoders before vs. after suppression of APC feedback. Positive values indicate higher decoding performance in the intact circuit.

#### Extended Data Table 1

| Odor Index | Odor Set A | Odor Set B |
|---|---|---|
| 1 | Allyl tiglate | 2,4 decadienal |
| 2 | Isoamyl acetate | valeraldehyde |
| 3 | Valeraldhyde | 2,3-Pentanedione |
| 4 | Ethyl Valerate | Ethyl hexanoate |
| 5 | Heptanal | Allyl butyrate |
| 6 | | Ethyl valerate |
| 7 | | 2,3-Diethylpyrazine |
| 8 | | Hexanal |
| 9 | | Ethyl heptanoate |
| 10 | | Heptanal |
| 11 | | Allyl tiglate |
| 12 | | ethyl tiglate |
| 13 | | Isoamyl acetate |
| 14 | | Methyl tiglate |
| 15 | | Cineole |
| 16 | | 2-hexanone |
| 17 | | isobutyl propionate |
| 18 | | Hexanoic acid |
| 19 | | 1,3 dimethoxybenzene |
| 20 | | Valeric acid |

#### References:


1. Shepherd, G. M. Synaptic organization of the mammalian olfactory bulb. *Physiol. Rev.* **52**, 864–917 (1972).
2. Shipley, M. T. & Adamek, G. D. The connections of the mouse olfactory bulb: a study using orthograde and retrograde transport of wheat germ agglutinin conjugated to horseradish peroxidase. *Brain Res. Bull.* **12**, 669–688 (1984).
3. Oswald, A.-M. & Urban, N. N. There and Back Again: The Corticobulbar Loop. *Neuron* **76**, 1045–1047 (2012).
4. Rothermel, M. & Wachowiak, M. Functional imaging of cortical feedback projections to the olfactory bulb. *Front. Neural Circuits* **8**, 73 (2014).
5. Boyd, A. M., Sturgill, J. F., Poo, C. & Isaacson, J. S. Cortical feedback control of olfactory bulb circuits. *Neuron* **76**, 1161–1174 (2012).
6. Boyd, A. M., Kato, H. K., Komiyama, T. & Isaacson, J. S. Broadcasting of cortical activity to the olfactory bulb. *Cell Rep.* **10**, 1032–1039 (2015).
7. Otazu, G. H., Chae, H., Davis, M. B. & Albeanu, D. F. Cortical Feedback Decorrelates Olfactory Bulb Output in Awake Mice. *Neuron* **86**, 1461–1477 (2015).
